# Interactome-based deep learning and path-level decomposition reveal TGFβ-IFNα competition in MASH

**DOI:** 10.64898/2026.10.24.761530

**Authors:** Nikolaos Meimetis, Tyler Matsuzaki, Xingrui Mou, Lindsey Foster, Linda G. Griffith, Douglas A. Lauffenburger

## Abstract

Dysregulated signaling drives Metabolic Dysfunction-Associated Steatohepatitis (MASH) through a complex crosstalk between pro- and anti-fibrotic cues. We previously observed potential competition between IFNα-mediated JAK-STAT signaling and TGFβ in modulating MASH fibrosis, but the precise mechanism of that competition, and whether it can act directly on hepatocytes, is yet unknown. We used DT-LEMBAS, an interactome-based deep learning framework, training an ensemble of 50 models on drug-induced transcriptomics from primary human hepatocytes. The ensemble captured transcriptional changes induced by simulated receptor knockouts and aligned predicted profiles with known TGFβ- and IFNα-induced phenotypes. Systematic simulations demonstrate their competitive modulation of MASH phenotype scores benchmarked against human cohorts. To extract mechanistic insights, we developed a gradient-based approach, where we computed the Jacobian of the used recurrent signaling network and applied chain rule decomposition to identify a four-node module of SMAD3, TP63, STAT2, and STAT3. Mechanistic hypotheses involving these canonical and non-canonical TF mediators were experimentally validated in primary hepatocyte spheroids. This work establishes a generalizable and scalable approach to decode large signaling neural networks into interpretable biological subnetworks.

## 1. Introduction

Dysregulated signaling, originating from multiple conditions such as genetic mutations, chemical stimuli, or environmental factors, underlies numerous diseases (Hanahan & Weinberg, 2011; Hanahan, 2022; García-Velázquez & Arias, 2017; Popugaeva *et al*, 2018; Kok *et al*, 2020; Deng *et al*, 2024). In metabolic dysfunction-associated steatotic liver disease (MASLD), for example, fibrosis progression emerges from a complex cross-talk between pro- and anti-fibrotic cues (Vachliotis & Polyzos, 2025; Xu *et al*, 2022; Ramachandran & Iredale, 2012; Steinberg *et al*, 2025).

This disorder is primarily associated with excessive caloric consumption and high-fat diets, alongside a sedentary lifestyle (Pafili & Roden, 2021), leading to the accumulation of excess fat in the liver (Geng *et al*, 2021). The global prevalence of MASLD is alarmingly high and continues to rise in parallel with the obesity epidemic, with the number of affected adults worldwide having increased from 25% between 1990 and 2006 to 38% in 2016-2019 (Wong *et al*, 2023). If left untreated, MASLD can progress into metabolic dysfunction-associated steatohepatitis (MASH), characterized by increased liver inflammation and fibrosis. These structural damages lead to an increased risk of many other comorbidities, including hepatocarcinoma, liver failure, and cardiovascular disease (Steinberg *et al*, 2025; Anstee *et al*, 2019; Mollace *et al*, 2024). The progression of MASLD is governed by a complex network of cellular signaling, while dysregulation of the hepatic signaling microenvironment plays an important role in the chronic inflammation, insulin resistance, and fibrosis associated with this disease (Xu *et al*, 2022; Pafili & Roden, 2021).

Within this signaling landscape, evidence has recently emerged that TGFβ and IFNα may be opposing regulators of MASH progression (Cadavid *et al*, 2026). In MASLD, TGFβ has been established as a master regulator of hepatic stellate cell (HSC) activation and extracellular matrix (ECM) deposition, directly leading to fibrotic phenotypes and contributing to MASLD progression (Yang *et al*, 2014; Dooley & ten Dijke, 2012). In contrast, IFNα is known to have anti-fibrotic effects in mice and can ameliorate steatosis in hepatocytes in mouse models (Inagaki *et al*, 2003; Wieser *et al*, 2018), while anti-fibrotic effects of interferons in general have been reported in other biological contexts too(Hasaneen *et al*, 2015; Oldroyd *et al*, 1999; Weng *et al*, 2007; Tuechler *et al*, 2025). From the literature, it is known that TGFβ may indirectly affect JAK-STAT signaling or induce a phenotype similar to that of JAK-STAT inhibition, specifically in the context of MAFLD. The inhibition of growth and proliferation, and the apoptotic phenotype of hepatocytes under TGFβ stimulation, induces a transcriptional response resembling JAK-STAT signaling inhibition (Deng *et al*, 2024; Dooley & ten Dijke, 2012; Romero-Gallo *et al*, 2005; Rawlings *et al*, 2004), while there is evidence supporting a possible direct antagonization by TGFβ of interleukin-induced JAK-STAT signaling (Bright & Sriram, 1998; Biffi *et al*, 2019). Systems-level analysis of MAFLD in vitro models has provided computational evidence of a potential competition between IFNα-mediated JAK-STAT signaling and TGFβ signaling during MAFLD fibrosis progression (Cadavid *et al*, 2026), while IFNγ-mediated JAK-STAT signaling can inhibit TGFβ signaling in a genetically modified and simplified cell system (Ulloa *et al*, 1999). Yet manually delineating this relationship across the full genome-scale signaling network in hepatocytes remains challenging.

Computational modeling, with the advent of Machine Learning (ML) and AI, can aid in the delineation of such complex signaling systems. Quite a few models exist nowadays that utilize genome-scale signaling networks to make predictions regarding cellular responses to external stimuli in an interpretable way (Nilsson *et al*, 2025). For example, Fortelny et al. used prior knowledge interaction networks to predict receptor stimulation from downstream observed transcriptomics. Meanwhile, the LEMBAS family of models uses a top-down approach where recurrent neural networks are primed with a prior knowledge network of intracellular molecular interactions to predict transcription factor (TF) activities from upstream nodes’ stimulation (Nilsson *et al*, 2022; Meimetis *et al*, 2024; Nordenstorm *et al*, 2026; Antonopoulos *et al*, 2026). However, while these neural networks excel at capturing non-linear signaling dynamics and provide an interpretable scaffold to study them, they still rely on completely heuristic approaches to prune down the larger network to actionable explanations of the mechanism of action of perturbations. Thus, a mathematically principled method is still not established for extracting compact, interpretable signaling pathways from trained LEMBAS-like models. This is exactly what is needed to fully comprehend competitive signaling effects such as that between IFNα and TGFβ.

In this study, we address both the deconvolution of TGFβ-IFNα signaling competition in hepatocytes and the establishment of a rational way to extract small mechanistic explanations from genome-scale ANNs of intracellular signaling. Here, we train an ensemble of 50 DT-LEMBAS (Meimetis *et al*, 2024) models, per time point, on transcriptional data of drugs tested on primary human hepatocytes (Uehara *et al*, 2010; Igarashi *et al*, 2015), yielding hepatocyte-specific signaling models for in silico experimentation (which is the ultimate goal of this study, not drug response predictions). DT-LEMBAS is an extension of the LEMBAS family of models that utilizes ensembles of artificial neural networks primed with prior knowledge of molecular interactions and known drug-target interactions to automatically infer drug-target interactions and their downstream effects on intracellular signaling (Meimetis *et al*, 2024). Leveraging this deep learning ensemble, we systematically simulated IFNα- and TGFβ-mediated signaling in hepatocytes to investigate their competitive interplay in the progression of MASH. The model accurately captures transcriptional changes induced by simulated knockouts of IFNα and TGFβ receptors, including downstream marker transcription factors (e.g., SMADs, IRFs, STATs), and aligns predicted profiles with TGFβ- and IFNα-induced phenotypes. Our simulations reveal a competition between TGFβ and IFNα that modulates fibrotic responses and identifies candidate perturbations for amplifying or disrupting this crosstalk. To interpret this genome-scale signaling network, we develop a path-level decomposition method, using Jacobian gradient-based edge pruning and decomposing the gradient flow inside the trained network based on the chain rule. This ensemble approach converges in the end on a compact regulatory module of four transcription factors: SMAD3, TP63, STAT2, and STAT3. We generate mechanistic hypotheses involving both canonical (SMAD3, STATs) and non-canonical (TP63) mediators of MASH-related TGFβ and IFNα signaling, and experimentally validate these predictions in primary human hepatocyte spheroids. Altogether, this study reveals the prevalent way of competition between IFNα and TGFβ in the context of MASH, and indicates potential ways for modulating the progression of the disease. Finally, it provides a generalizable strategy for decomposing genome-scale signaling neural networks into biologically interpretable subnetworks.

## 2. Results

### 2.1. A hepatocyte-specific DT-LEMBAS ensemble accurately predicts drug-induced transcription factor activity in primary human hepatocytes

The relationship between interferon-mediated signaling and TGFβ signaling has so far been studied mostly in reduced systems, revealing evidence of competition in the case of IFNγ (Ulloa *et al*, 1999), with some corroboration of such evidence in liver cells of mice and rats (Park *et al*, 2025; Weng *et al*, 2007). These preclinical models, however, have substantial differences from human patients in liver diseases (Teufel *et al*, 2016; Vacca *et al*, 2024; Santhekadur *et al*, 2018). In human hepatocytes, the transcriptional response to TGFβ stimulation resembles an inhibition of JAK-STAT signaling (which interferons can in turn modulate), and there is limited evidence of direct antagonism between the two pathways in other cellular and disease contexts (Bright & Sriram, 1998; Biffi *et al*, 2019). We recently validated the direct antagonism, both phenotypically and in terms of transcriptomics (Cadavid *et al*, 2026). However, the exact signaling interplay between IFNα and TGFβ in human hepatocytes is not yet known, and to effectively take therapeutic advantage of this relationship, we would need to study signaling at the genome scale.

To obtain a genome-scale signaling model of human hepatocytes, we trained an ensemble of 50 independent DT-LEMBAS models (Meimetis *et al*, 2024) on microarray gene expression data from the Japanese Toxicogenomics Project (Igarashi *et al*, 2015; Uehara *et al*, 2010), which profiles transcriptional responses of primary human hepatocytes to 120 unique drugs across three doses and three time points of exposure (Fig. 1A). A different ensemble of 50 models was trained for each time point (2, 8 and 24 hours), resulting in 150 models in total. DT-LEMBAS models were trained to predict transcription factor (TF) activity, which was calculated (see more in section 4.1.2) using the VIPER algorithm (Alvarez *et al*, 2016a) coupled with the DoRothEA regulon (Garcia-Alonso *et al*, 2019). It is important to note, however, that our goal is not to construct a model predicting the transcriptional response of hepatocytes under drug perturbations, but rather to use the signaling part to perform in silico simulation of TGFβ and IFNα stimulations. However, first we need to establish that the model’s signaling network can be trusted via the predictions we can make in this specific dataset. The main benchmarking results presented here are from the 8-hours models, and the corresponding results for the 2- and 24-hours models are present in Appendix Fig. S1-2.

**Figure 1:**
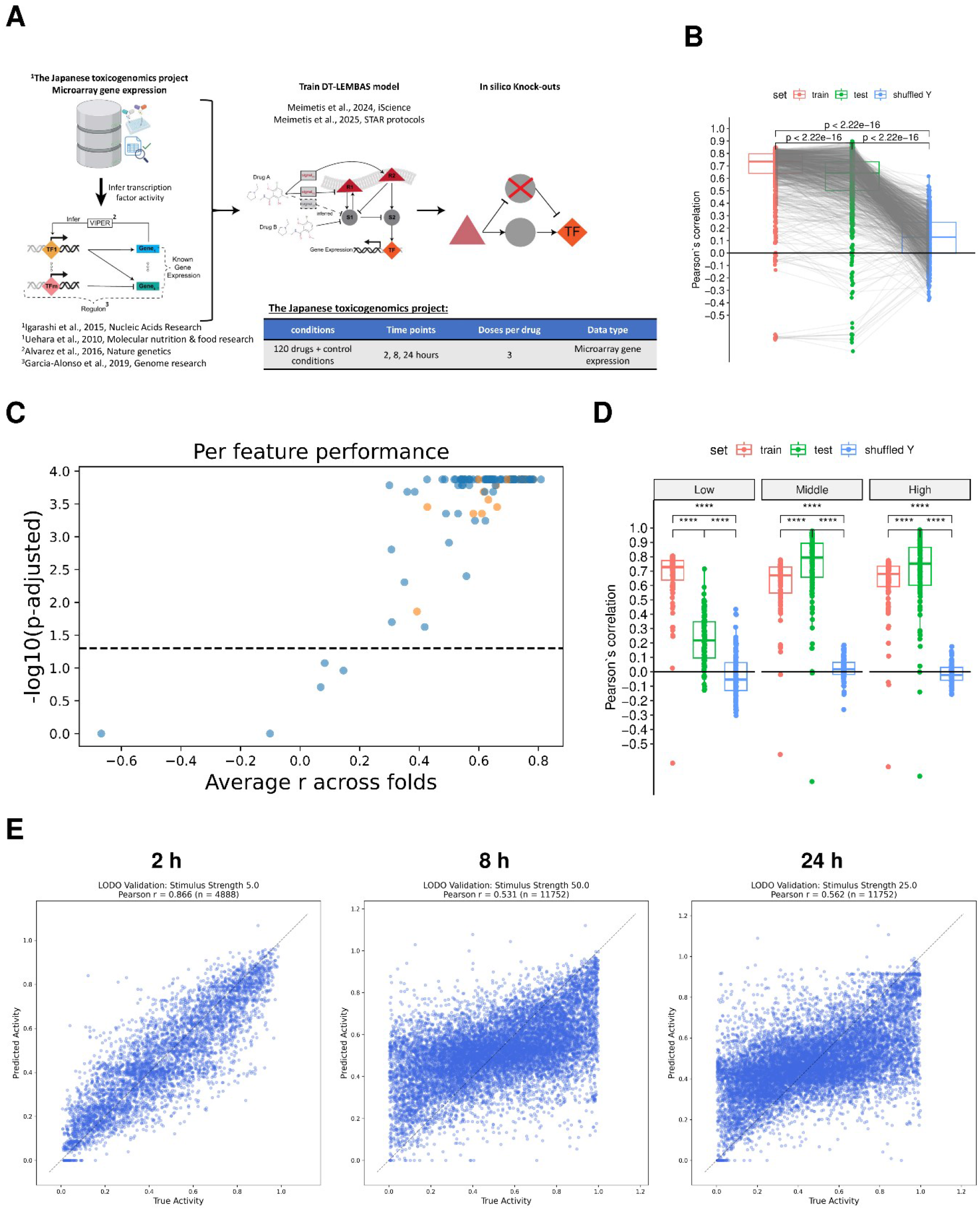
Training and predictive performance of hepatocyte-specific DT-LEMBAS models. **A)** Workflow for constructing hepatocyte signaling models and simulating perturbations (see section 4.2 for details). **B)** Predictive performance at 8 hours in random 10-fold cross-validation (CV). Each point represents the Pearson correlation between predicted and observed TF activity for one TF in one of the folds. **C)** Test performance for individual TFs at 8 hours. Each point represents one TF, with its mean test set Pearson correlation across folds plotted against the BH-adjusted p-value from a one-sided Mann-Whitney U test comparing test and randomized correlations. Orange points denote TFs previously associated with MAS or fibrosis stage. **D)** Leave-One-dose-Out cross-validation (LOOCV) at 8 hours. Each point represents the Pearson correlation for one TF in the indicated dose split, comparing training, held-out test and shuffled Y predictions. **E)** Leave-one-drug-out (LODO) validation at 2, 8 and 24 hours. Each point represents one drug-TF pair. Dashed diagonal lines denote perfect agreement. Comparisons in B and D used paired two-sided Wilcoxon signed-rank tests. In the boxplots, center lines denote medians, box bounds denote the first and third quartiles, and whiskers extend to the most extreme values within 1.5 times the interquartile range from the box. ****p-value≤ 0.0001.

The ensemble’s performance in predicting drug-induced TF activity was validated in two complementary ways. First (Fig. 2A), its accuracy in predicting TF activity was benchmarked against randomized models derived from shuffling the TF labels (“shuffled Y”) in a random 10-fold cross-validation (CV) scheme (Fig. 1B-C, Appendix Fig. S1-2). This provides a quantitative, TF-by-TF estimate of which TFs can be reliably predicted better than random (Fig. 1B). The model outperformed random expectation for 99 of 104 TFs, including all TFs previously found to be important for predicting MAS and fibrosis stage in human clinical data (Cadavid *et al*, 2026), based on a Partial Least Squares Regression (PLSR) model (Appendix Fig. S3). Secondly (Fig. 2B), a Leave-One-Out Cross-Validation (LOOCV) scheme was constructed in which, for each drug, the high, middle, or low dose was withheld from training in turn, to test whether the model could extrapolate to unseen doses (Fig. 1C). This scheme was likewise benchmarked against shuffled Y baselines. The model also outperformed the random baseline in this LOOCV scheme, indicating that it can extrapolate to relatively unseen drug doses (Fig. 1D). Additionally, even though the goal is not to use the model for drug response prediction, we evaluated how our model compares to a simpler linear model (extrapolating or interpolating between doses for the activity of each TF). Across time points and dosing schemes, neither architecture was uniformly superior. Instead, the relative performance depended on exposure time, dose, and TF (Fig. 3). We therefore use the linear comparison to identify where the network-constrained nonlinear architecture adds predictive information, rather than to claim blanket superiority, as it may be the case that a linear assumption for dosing schemes is enough to explain the data. Nevertheless, the ensemble of models has a slight edge over the linear baseline (Fig. 3), further validating that the models have learned useful information.

**Figure 2:**
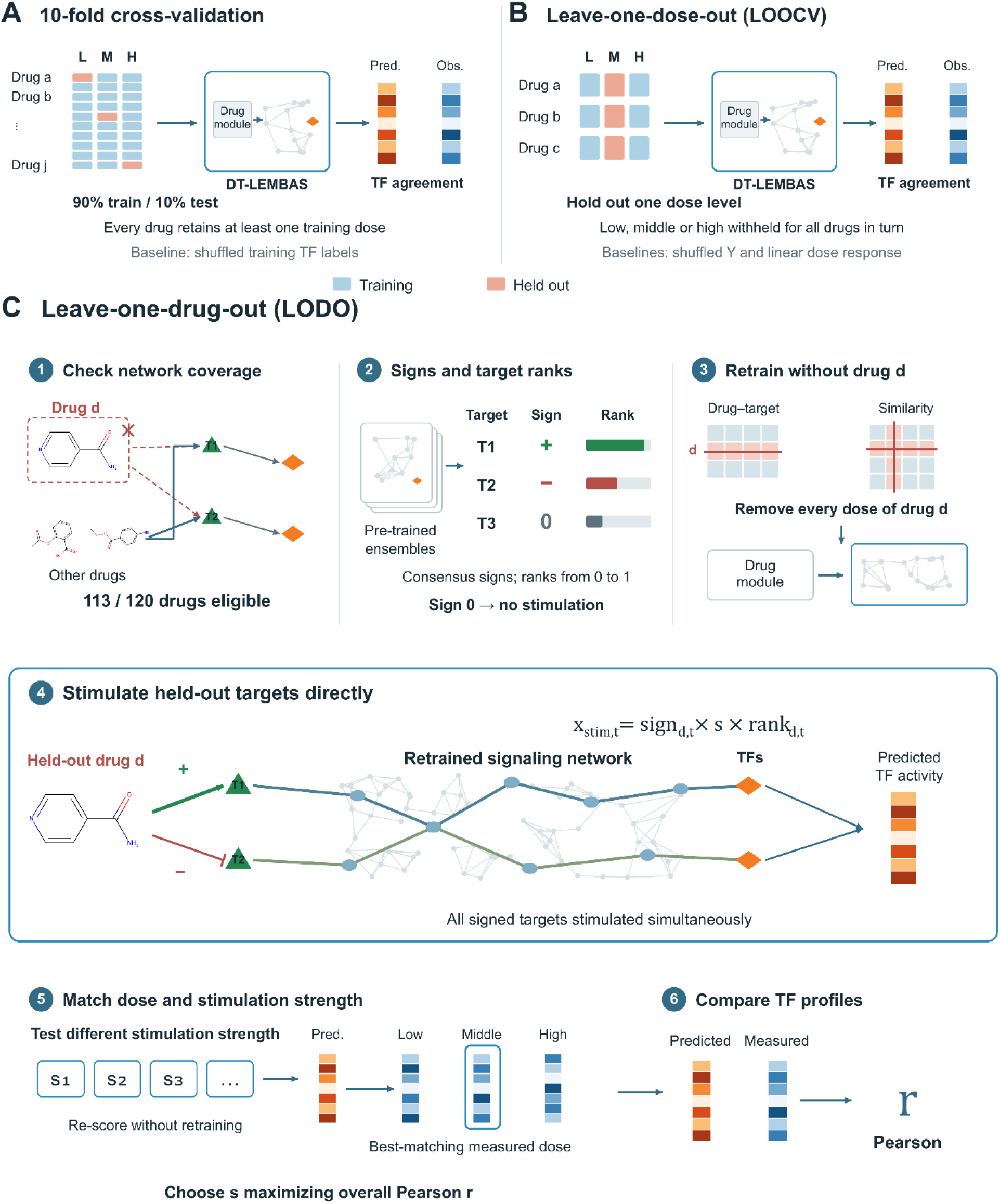
Validation procedures used to evaluate drug-induced TF activity predictions. **A)** Random 10-fold cross-validation (CV). In each split, 90% of the drug-dose profiles were used for training and 10% for testing, with at least one dose of every drug retained in training. Predictions were compared with a model retrained using shuffled training TF labels. **B)** Leave-one-dose-out cross-validation (LOOCV). The low, middle or high dose was withheld for all drugs in turn. Predictions were compared with shuffled Y models and with linear interpolation or extrapolation fitted separately for each drug and TF. **C)** LODO validation procedure (see section 4.2.2.3). Network coverage was first checked to identify drugs whose targets remained represented by, or reachable from, the perturbations retained during training. Consensus interaction signs and target ranks were obtained from the pre-trained ensembles. For each held-out drug, all of its doses and its entries in the drug-target and chemical similarity matrices were removed, and one model was retrained per time point using the same signaling network. All targets with non-zero consensus signs were then stimulated simultaneously, bypassing the drug module. Agreement was evaluated per TF, per drug and across all drug-TF pairs. Each validation was performed separately at 2, 8 and 24 hours.

**Figure 3:**
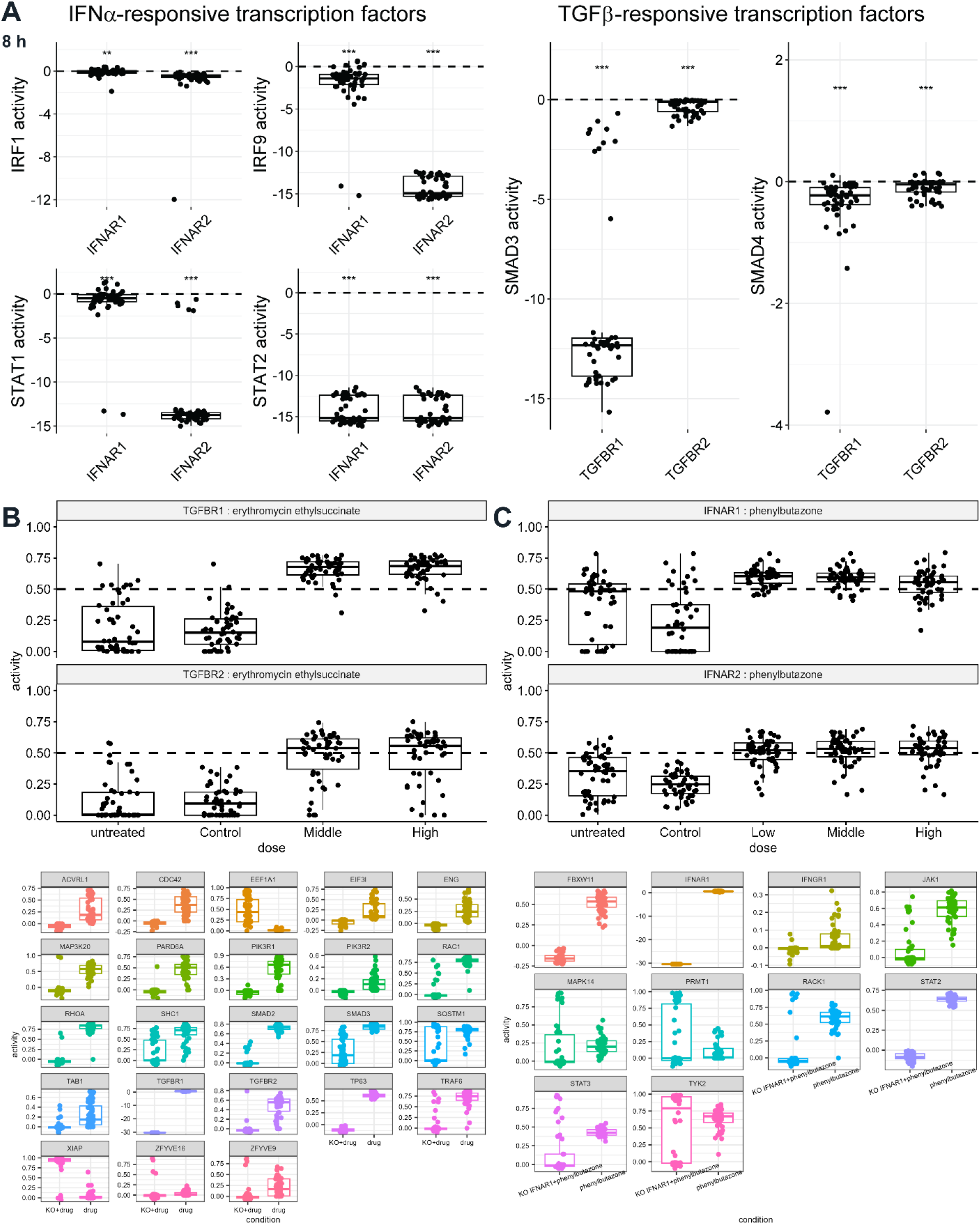
Receptor perturbations recover expected signaling responses and identify drug conditions that engage TGFβ and IFNα receptors. **A)** Change in the predicted activity of IFNα-responsive TFs (IRF1, IRF9, STAT1 and STAT2) following IFNAR1 or IFNAR2 knockout, and of TGFβ-responsive TFs (SMAD3 and SMAD4) following TGFBR1 or TGFBR2 knockout at 8 hours. Changes were calculated relative to each model’s untreated state. The dashed line denotes no change. p-values were obtained from one-sided one-sample Wilcoxon signed-rank tests against zero, testing for a decrease in activity. **B)** Predicted TGFBR1 and TGFBR2 activities under untreated, matched-control, middle-dose and high-dose erythromycin ethylsuccinate conditions (upper panels), and predicted activities of the indicated signaling nodes under erythromycin ethylsuccinate with or without TGFBR1 knockout (lower panels). **C)** Predicted IFNAR1 and IFNAR2 activities under untreated, matched-control, low-dose, middle-dose and high-dose phenylbutazone conditions (upper panels), and predicted activities of the indicated signaling nodes under phenylbutazone with or without IFNAR1 knockout (lower panels). All panels show simulations from the 8-hour ensemble. Each point represents one of 50 independently trained models. In all boxplots, center lines denote medians, box bounds denote the first and third quartiles, and whiskers extend to the most extreme values within 1.5 times the interquartile range from the box. *p-value< 0.05, ** p-value < 0.01 and *** p-value < 0.001.

Because only the signaling network component of DT-LEMBAS (not the drug module) is used in the simulations that follow, its ability to reproduce in silico receptor knockout and knockdown behavior was validated separately. Apart from the response of known downstream markers of IFNα and TGFβ, a systematic evaluation of the stimulation of signaling nodes in silico needs to be performed. On this front, we performed a Leave-One-Drug-Out (LODO) validation, where a whole drug is kept out of training, and then its own targets are stimulated in silico (Fig. 2C, Fig. 1E). The TF activity induced by the in silico stimulation and the known drug-induced TF activity are compared. To avoid network coverage artifacts, this analysis was restricted to drugs whose targets remained directly represented in, or reachable from, the perturbations retained during training. For each eligible drug, the model was retrained without that drug, after which all of its annotated targets with a non-zero consensus sign were perturbed simultaneously. The sign, determined by majority consensus across the previous 2, 8, and 24 hour ensembles, specified activation or inhibition. Perturbation magnitudes were target-specific rather than uniform (see 4.2.2.3), i.e., a common base strength was multiplied by a consensus target rank (in the range of 0 to 1) derived from the learned drug-target interaction scores, such that higher-ranked targets received stronger inputs. Because this artificial stimulation does not correspond to a physical drug concentration, multiple base strengths were evaluated. Identifying the exact value of the targets’ artificial stimulation accounting for biophysical properties of the known interactions (e.g. binding affinity) and the drug concentration is beyond the scope of this work. Since this in silico stimulation is dose-agnostic, for every drug we compared the predicted TF activity against the experimental dose whose measured profile it best matched, quantifying agreement by Pearson correlation. We further used a range of base stimulation strengths and retained the value that maximized the overall correlation. The strength maximizing the overall Pearson correlation across the selected drug–TF pairs was retained separately for each time point (see 4.2.2.3). Thus, this validation tests whether the signaling network can reconstruct the transcriptional response to a completely unseen in silico perturbation, from signaling node-level stimulation alone. The models recovered the drug-induced TF activity (Fig. 1E), indicating that the signaling network itself, and not merely the drug module, learned a meaningful input-to-TF mapping. Finally, this performance is a preliminary indication for future work on in silico stimulations of drugs’ targets not initially included in the models’ drug space.

### 2.2. Simulated knockouts of TGFβ and IFNα receptors recapitulate established downstream signaling and align with clinical transcriptional patterns

After validating the ability of the models to make predictions on out-of-distribution data (establishing that the learnt structure is trustworthy), we reasoned that a hepatocyte signaling model suitable for in silico experimentation should reproduce the immediate transcriptional consequences of perturbing canonical cytokine receptors. We therefore simulated loss of TGFBR1, TGFBR2, IFNAR1, and IFNAR2 by assigning a large negative value as its input, and quantified the resulting changes in TF activity across the corresponding 50 models’ ensemble. Knockout of IFNAR1 or IFNAR2 decreased the activity of established interferon-responsive factors, including IRF1, IRF9, STAT1, and STAT2. Conversely, knockout of TGFBR1 or TGFBR2 reduced SMAD3 and SMAD4 activity (Fig. 3A). Because the knockdown is induced with a large negative input, we confirmed that at the chosen value the activities of neighboring input and output nodes, of all predicted TFs, and of the resulting phenotype scores had converged (Appendix Fig. S4-6).

We then examined whether the receptor knockdowns turned off (or on, depending on how the receptor affects downstream nodes) the activity of the receptors and their downstream nodes when the network is already stimulated with identified activators of the receptors from our training data. At 8 hours, erythromycin ethylsuccinate was associated with higher predicted TGFBR1 and TGFBR2 activities at the displayed middle and high doses than in the untreated and matched-control states (Fig. 3B). Phenylbutazone similarly increased the predicted activities of IFNAR1 and IFNAR2 across the displayed dose range (Fig. 3C). These examples show that the ensembles can map drug-induced transcriptional states onto coherent receptor programs. Similar results can be observed using the 2- and 24-hours models (Appendix Fig. S7-8).

Finally, to connect the simulated signaling states to human disease, we projected the predicted TF activities onto MAS and fibrosis stage axes derived from patient transcriptomic data. A PLSR model was used to translate TF activities into phenotype scores. At 8 hours, TGFBR1 knockout decreased both MAS and fibrosis stage scores, whereas IFNAR1 and IFNAR2 knockout increased both scores (Fig. 4A). TGFBR2 knockout produced a smaller shift that remained closer to the untreated distribution, but this may be an artifact of the prior knowledge network rather than true biology.

**Figure 4:**
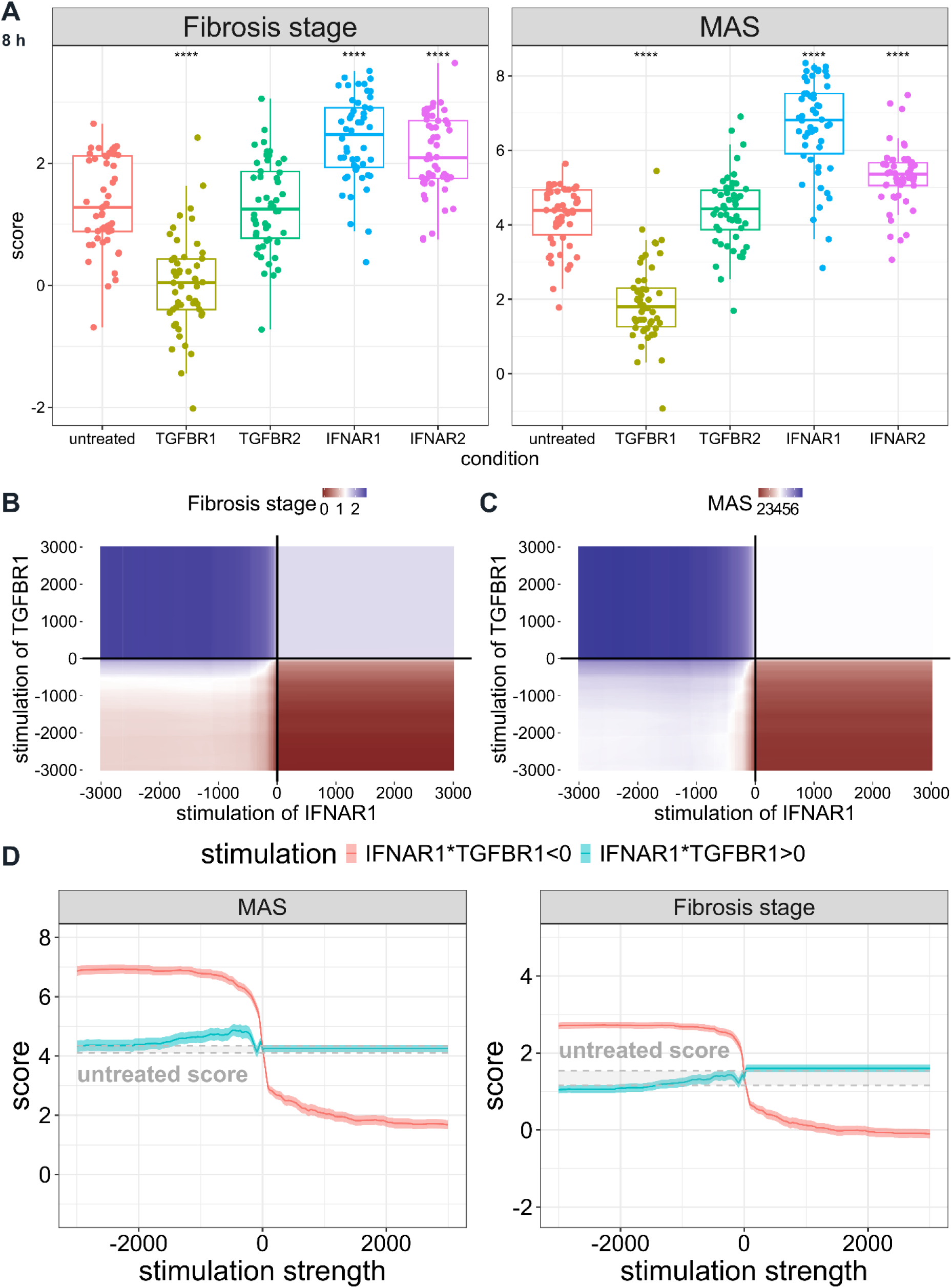
TGFBR1 and IFNAR1 exert opposing control over predicted MAS and fibrosis stage. **A)** MAS and fibrosis stage scores obtained by applying a partial least-squares regression model (trained on human liver transcriptomic data) to the TF activities predicted after receptor knockout at 8 hours. Each point represents one of 50 independently trained models. Receptor-knockout conditions were compared with the untreated condition using two-sided Wilcoxon rank-sum tests. Boxplots show medians and first and third quartiles, with whiskers extending to the most extreme values within 1.5 times the interquartile range from the box. ****p-value ≤ 0.0001. **B, C)** Ensemble-mean fibrosis stage (B) and MAS (C) scores across the two-dimensional TGFBR1-IFNAR1 stimulation grid at 8 hours. The color scales are centered on the corresponding untreated scores, with blue indicating higher scores and red indicating lower scores. **D)** Predicted MAS and fibrosis stage scores along the opposing-sign diagonal (IFNAR1 × TGFBR1 < 0, coral color) and the co-directional diagonal (IFNAR1 × TGFBR1 > 0, turquoise color) of the stimulation grid. The horizontal axis denotes IFNAR1 input. TGFBR1 receives an input of equal magnitude with the opposite or the same sign, respectively. Lines show means across 50 models and colored ribbons show ±1 standard error (SE) of the mean. Grey shading and dashed lines indicate the untreated reference. Receptor stimulation strengths are expressed in arbitrary simulation units, with negative values representing inhibition and positive values representing activation.

The direction of these phenotype changes is consistent with pro-fibrotic TGFβ responses and anti-fibrotic IFNα responses from various disease contexts(Dooley & ten Dijke, 2012; Deng *et al*, 2024; Tuechler *et al*, 2025; Park *et al*, 2025). Importantly, the phenotype projection is trained on human clinical variation, whereas the receptor perturbations are generated within the hepatocyte-specific signaling network. Their agreement links the receptor-level simulations to disease-associated transcriptional patterns without assuming that the two datasets contain matched perturbations.

### 2.3. TGFβ and IFNα competitively modulate MASH fibrosis and steatohepatitis phenotypes in silico

Having established the opposing effects of receptor loss, we next asked how the two signals interact when their strengths are varied simultaneously. We simulated a 2-dimensional (2D) grid of TGFBR1 and IFNAR1 perturbations, calculated the resulting TF activities for each model at 8 hours, and converted these activities to MAS and fibrosis stage scores using the same PLSR model. The response surfaces were organized primarily by the relative direction of the two receptor inputs rather than by either input in isolation (Fig. 4B–C).

When TGFBR1 and IFNAR1 were perturbed in opposite directions, their effects reinforced each other. TGFBR1 activation together with IFNAR1 inhibition produced the highest MAS and fibrosis stage scores, whereas TGFBR1 inhibition together with IFNAR1 activation produced the lowest scores. In contrast, co-directional perturbations of the two receptors remained closer to the untreated phenotype over a broad range of simulated input strengths. This relationship was apparent when the two diagonals of the stimulation grid were summarized as one-dimensional response curves (Fig. 4D). The same observations are made at 2-hour and 24-hour models (Appendix Fig. S9).

These results support a competitive relationship in which IFNα-associated signaling counterbalances the phenotype shift induced by TGFβ-associated signaling. The simulated input scale is an arbitrary model perturbation rather than a calibrated cytokine concentration. Accordingly, the grid identifies the direction and balance of the interaction but should not be interpreted as a biological dose-response curve.

Sequential simulations (see Appendix Supplementary Methods 1) further suggest that this balance depends on timing; i.e. introducing IFNAR1 stimulation after the TGFBR1-associated state was established did not reproduce the counterbalancing effect of simultaneous inputs (Appendix Fig. S10). We tested that this observation is not necessarily an artifact of accumulating errors of multiple models by evaluating that indeed the sequential simulation method can be used to correctly predict TF activities (Appendix Fig. S11, Appendix Supplementary Methods 1).

### 2.4. Jacobian-based gradient pruning identifies key receptor-to-phenotype signaling axes in the trained ensemble

The competitive response emerged from networks containing thousands of molecular interactions; however we identifying key drivers for experimental validation and further perturbation is not tractable using all thousands of interactions. Thus, we developed a path-level decomposition approach (Fig. 5A, Methods) that scores paths from selected receptors (here TGFBR1 and IFNAR1) towards a downstream node of interest (here MAS and fibrosis stage separately). By selecting the top 3 paths for each of the 50 models in an ensemble, we extract the subnetwork explaining mostly the effect of IFNAR1 and TGFBR1 on MAS and fibrosis, for each of the timepoints available. We first linked the phenotypes to the network with weights from a ridge regression predicting MAS and fibrosis stage from TF activity in the human cohort. Briefly, we scored each with an integrated-gradient approach, accumulating the single-step Jacobian of the recurrent network at equilibrium and multiplying along each path by the chain rule, integrated from a zero baseline to full stimulation (see section 4.4 for details). To avoid penalizing longer paths heavily, scores were normalized by a geometric mean along the path, and the top scoring paths per model were aggregated into an ensemble edge frequency score (Fig. 5A). Applied to the 2, 8, and 24 hour ensembles, the decomposition yielded compact, timepoint-specific subnetworks (Fig. 5B). Interestingly, while differences are indeed observed, the aggregated networks converged on four TFs that remained connected to the receptor inputs at all three time points (Fig. 5B). SMAD3, TP63, STAT2, and STAT3 were identified as the dominant mediators connecting the TGFβ and IFNα receptors to the MASLD phenotypes, and explaining their competitive effects. Additional intermediates, including MAPK14, FOXO4, and HIF1A, appeared in a time-dependent manner, but they did not replace the four-TF core.

**Figure 5:**
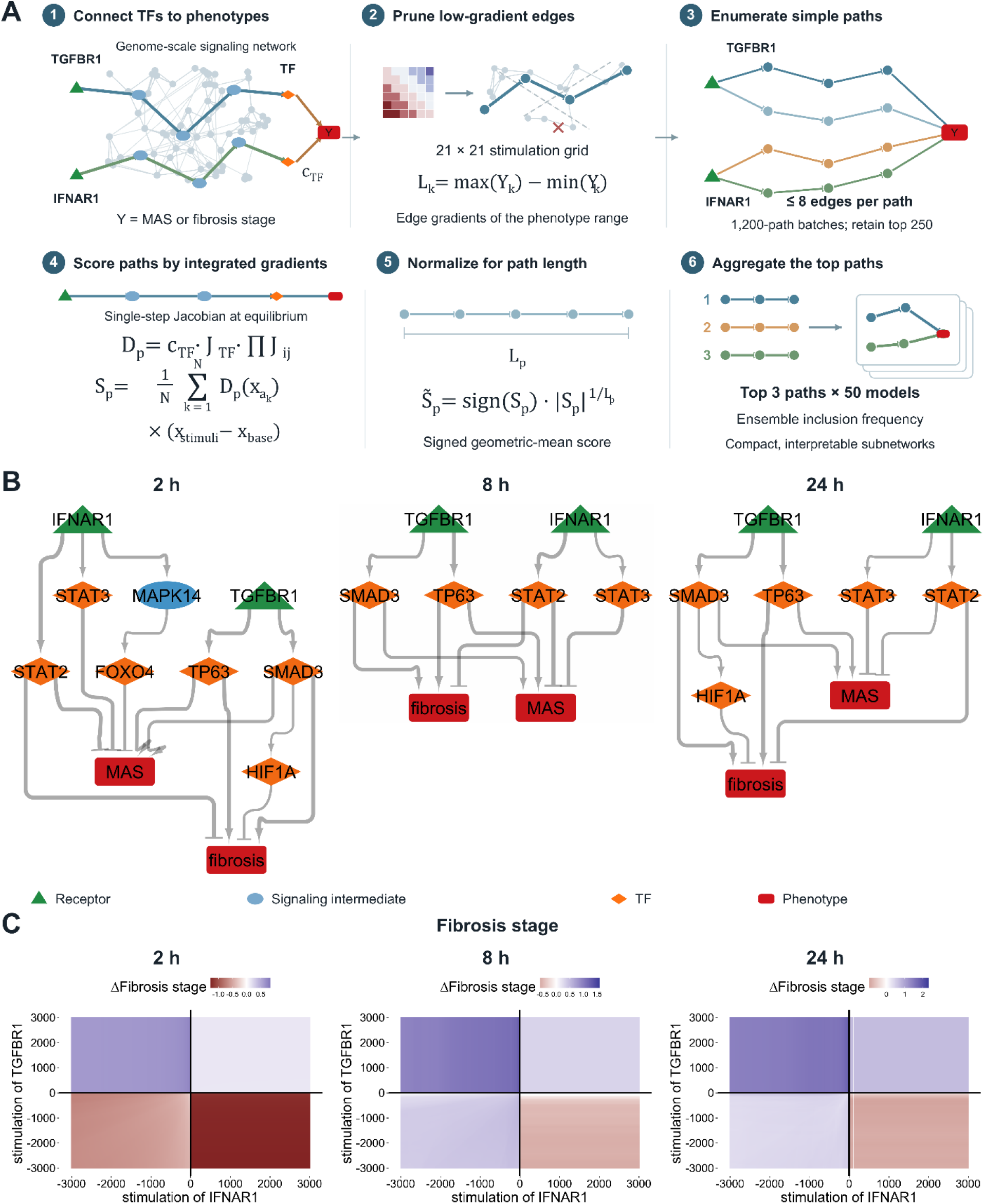
Jacobian and chain-rule decomposition identifies compact receptor-to-phenotype signaling networks. **A)** Procedure (see section 4.4 for more details) for decomposing genome-scale signaling to the receptor-to-phenotype paths, that mostly explained the mechanism of action of the stimulated receptors on the phenotypes of interest. D_p_, S_p_ and S̃_p_ denote the contribution at one integration step, the integrated path score and its length-normalized value, respectively. L_p_ is the number of edges in the path. The TF-to-phenotype coefficient is denoted by c_TF_, the TF output-scaling derivative by J_TF_, and the local signaling Jacobian by J_ij_. **B)** Subnetworks obtained for the 2-, 8- and 24-hours ensembles by combining paths linked to MAS and fibrosis stage. Receptors are shown as green triangles, signaling intermediates as blue ovals, TFs as orange diamonds and phenotypes as red rectangles. Arrowheads and blunt ends indicate positive and negative modeled interactions, respectively. **C)** Ensemble-mean changes in predicted fibrosis stage across the TGFBR1-IFNAR1 stimulation grid after restricting the signaling models to the extracted subnetworks at 2, 8 and 24 hours (similar to Fig. 4B).

SMAD3 is a well-known regulator of fibrosis (Deng *et al*, 2024; Xu *et al*, 2016), and SMAD3-knockout mice have been shown to be more resistant to developing obesity and insulin resistance on fatty diets (Tan *et al*, 2011). Tp63 is the gene that produces the transcription factor p63. While there is some evidence that the TAp63 isoform has a role in regulating steatosis in mouse hepatocytes (Porteiro *et al*, 2017; Fondevila *et al*, 2024), the influence of TGF-β on its function (or its effect directly on human hepatocytes) has not yet been reported. The remaining two transcription factors, STAT2 and STAT3, are downstream of the IFN-α receptor. Although relatively less studied, STAT2 is suggested to have a protective effect against fibrosis. STAT3 has been shown to have contradictory effects, with some studies showing it can reduce inflammation in MASH, whereas others demonstrate STAT3 can exacerbate fibrosis (Liu *et al*, 2023). Importantly, there has been limited investigation into the interplay of these four transcription factors in MASH signaling. However, their appearance across all three temporal networks suggests that they may play a key role in regulating MASH-related phenotypes.

The reduced subnetworks (Fig. 5B) reproduced the phenotypic behavior of the full model (Fig. 5C). Then, to test whether these TFs were functionally required within the simulated competition, we repeated the co-directional and opposite-direction TGFBR1/IFNAR1 perturbations after knocking out (in silico) SMAD3, STAT2, STAT3, TP53, or TP63 (Fig. 6A). The same panel was evaluated at 2, 8, and 24 hours. We additionally included JUN and MYC because they were considered alternative high-weight TFs in the phenotype projection (Appendix Fig. S12). The no-knockout panel reproduced the competitive separation we observed in the full model. STAT2 knockout produced the most specific loss of the IFNα anti-fibrotic (and anti-MAS) effect (Fig. 6B). The IFNAR1-driven reduction in MAS and fibrosis was diminished, leaving the TGFβ-associated phenotype comparatively less opposed. The TP63 knockout produced the reciprocal pattern, attenuating the pro-MAS and pro-fibrotic TGFβ-associated shift and narrowing the separation between the two input diagonals, particularly for MAS (Fig. 6C). These effects were qualitatively preserved across time, although their magnitude varied between the 2, 8, and 24 hours ensembles (Appendix Fig. S13-15), although we cannot evaluate if this is true biology with these exact data (they serve for hypothesis generation). SMAD3 knockout lowered the overall MAS and fibrosis stage scores but retained a larger separation between the input grid than TP63 knockout in several MAS simulations (Fig. 6D). STAT3 and TP53 knockout similarly shifted the phenotype level rather than selectively removing the antagonistic relationship (Fig. 6E). This distinction is important: a TF can strongly influence the phenotype score without being the node that most specifically couples the two receptor programs. Within the model, the knockout scan therefore supports STAT2 and TP63 as complementary mediators of the IFNα and TGFβ arms, while retaining SMAD3 as a major canonical determinant of the absolute disease-associated state.

**Figure 6:**
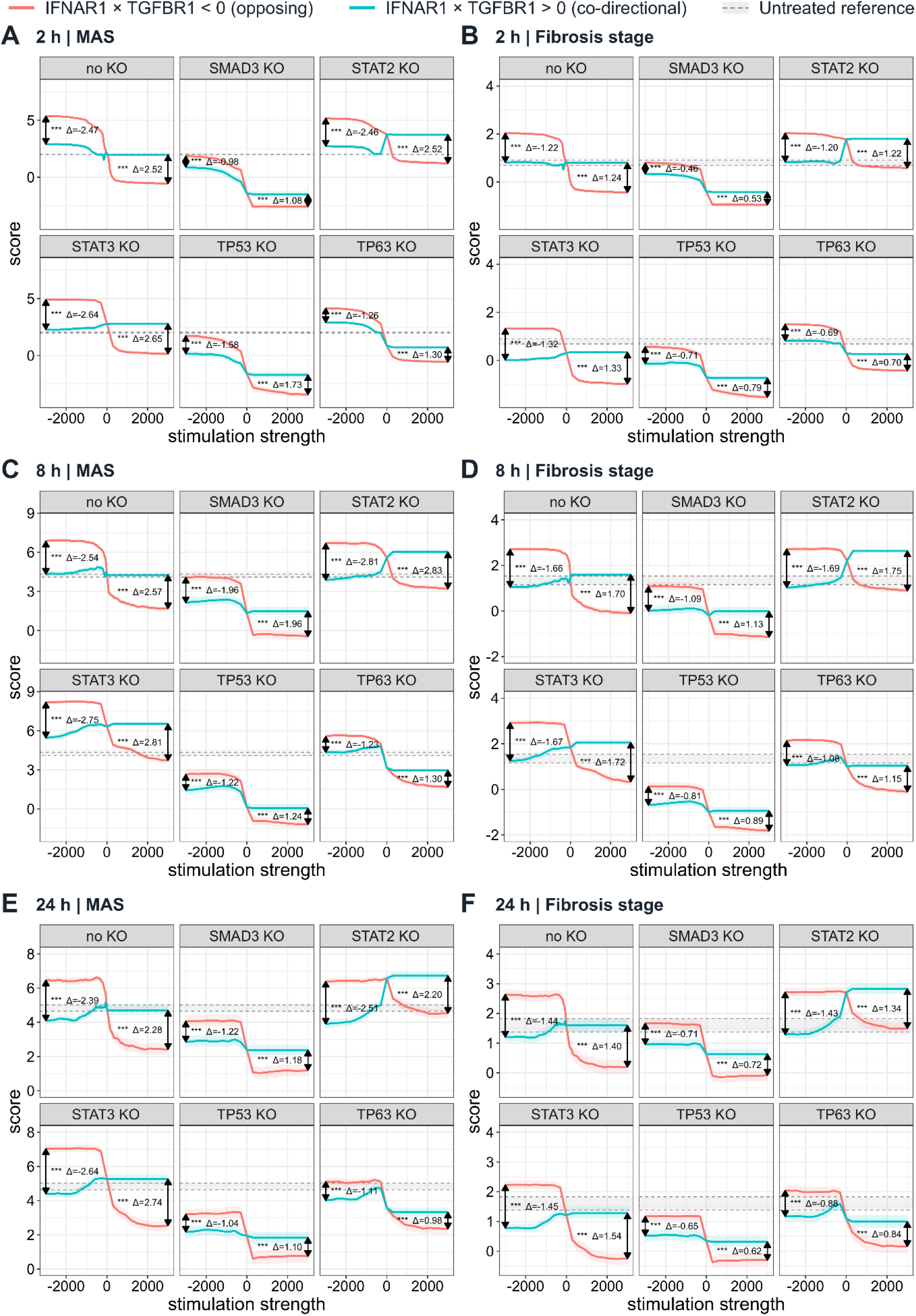
Mediator knockouts reshape the phenotypic response to opposing and co-directional receptor perturbations. A–F) Predicted MAS and fibrosis stage scores at 2 hours (A, B), 8 hours (C, D) and 24 hours (E, F), respectively, during combined TGFBR1 and IFNAR1 stimulation. Each panel shows the network without a mediator knockout (no KO) and after in silico knockout of SMAD3, STAT2, STAT3, TP53 or TP63. TF activities were converted to phenotype scores using the clinical partial least-squares regression model. Coral lines show opposing-sign receptor inputs (IFNAR1 × TGFBR1 < 0), and turquoise lines show co-directional inputs (IFNAR1 × TGFBR1 > 0). The horizontal axis denotes IFNAR1 input, with TGFBR1 receiving an input of equal magnitude and either the opposite or the same sign. Lines show means across the 50 models at each time point, and colored ribbons show ±1 SE of the mean. Grey shading and dashed lines indicate the untreated reference. Δ is the co-directional score minus the opposing-sign score. Comparisons at the extremes used paired two-sided t-tests, with Bonferroni adjustment across the comparisons within each phenotype and time point. *p-value < 0.05, ** p-value < 0.01 and *** p-value < 0.001.

Altogether, these data provide evidence that our Jacobian-based gradient pruning can reveal key transcription factors in MASH progression that may otherwise be hidden in complex neural networks and yield a compact set of transcriptional regulators whose functions can allow for direct experimental investigation. We corroborate these findings from the Jacobian-based method, with an orthogonal framework for pruning down the DT-LEMBAS model (Appendix Fig. S16). We thought that a Gaussian Graphical Model (GGM) could be another appropriate method for retrieving subnetworks using our simulated data (Appendix Supplementary Method 2), since GCMs have been successfully applied to model signaling interactions in microarray transcriptomic data (Dobra *et al*, 2004; Pihur *et al*, 2008).

### 2.5. RNA profiling in hepatocyte spheroids supports selected cytokine responses predicted by the model

To examine whether the model-derived responses could be observed experimentally, we profiled the transcriptome from primary human hepatocyte spheroids cultured in high-fat/type 2 diabetes (T2D) base medium following IFNα, TGFβ, or combined treatment. We inferred the activities of the four TFs prioritized by the network decomposition and examined disease-associated phenotype projections derived from gene expression and TF activity.

TGFβ-treated spheroids showed higher inferred SMAD3 and TP63 activities than the DMSO condition, whereas IFNα-treated spheroids showed higher STAT2 activity (Fig. 7A). STAT2 activity also remained high in the combined condition. These observations support the expected TGFβ–SMAD3 and IFNα–STAT2 responses and provide transcriptional evidence for the TGFβ-associated TP63 response nominated by the decomposition. However, inferred TP63 activity does not establish changes in p63 protein abundance, phosphorylation, or isoform-specific function.

**Figure 7:**
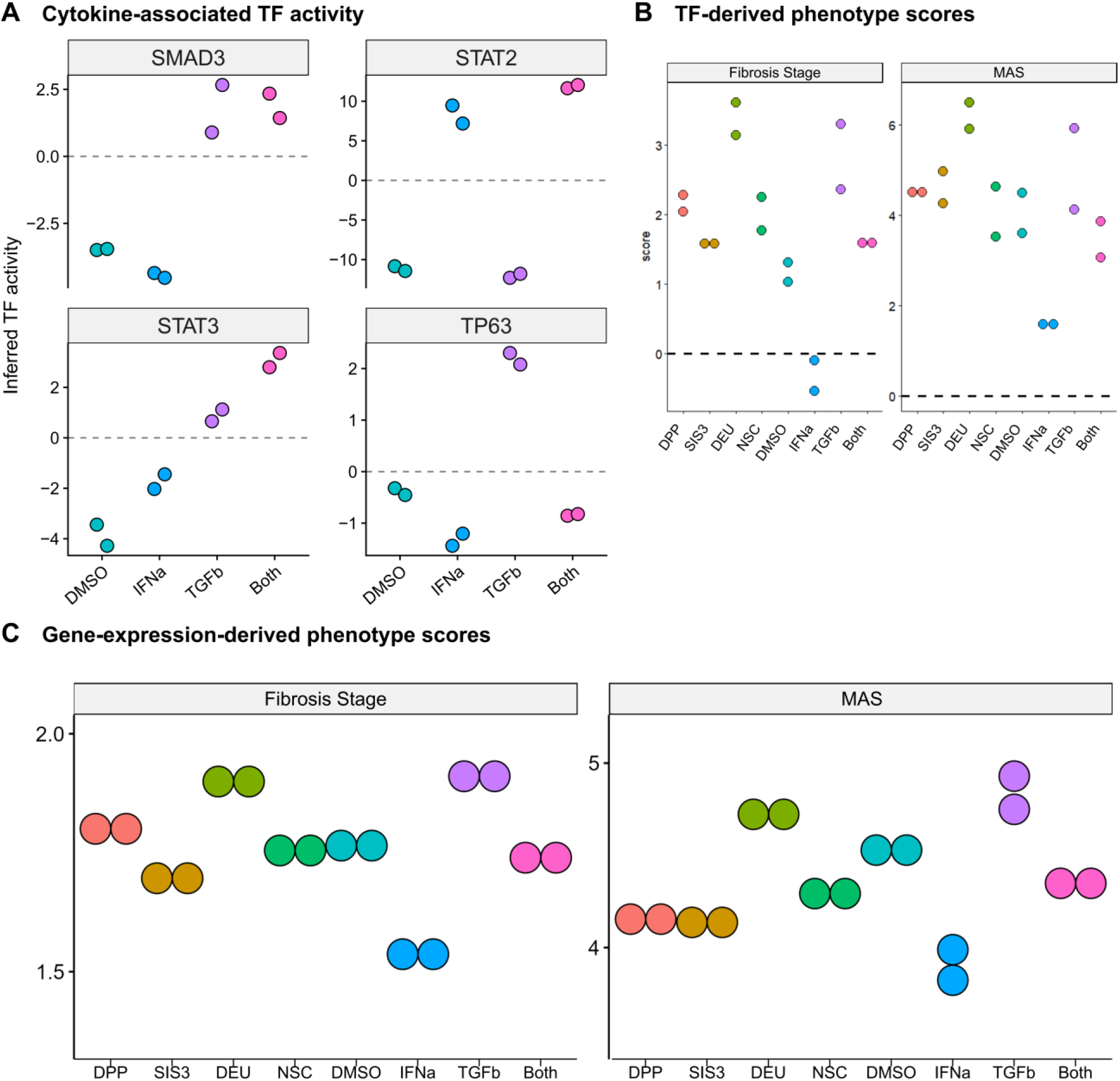
RNA profiling of hepatocyte spheroids supports selected model-derived cytokine responses. A) VIPER-inferred TF activities under DMSO, IFNα, TGFβ and combined treatment (Both). Points denote RNA profiles; offsets do not imply pairing. Dashed lines mark dataset-relative zero, not an untreated rteference. B, C) MAS and Fibrosis Stage projections from total TF activity (B) or gene expression (C); C uses shortened y-axis ranges. Compound labels are retained as assayed. Two RNA profiles are shown per condition, without inferential statistics. Scales vary across facets. Phenotype scores are continuous RNA-derived projections, not histological grades; negative scores do not imply negative biological fibrosis.

The combined condition showed different relationships for the individual TFs. SMAD3 activity remained high and overlapped the range observed with TGFβ alone, whereas TP63 activity remained inhibited, in contrast to the activation values under TGFβ alone (Fig. 7A). Thus, the combined condition retained the SMAD3 response associated with TGFβ and the STAT2 response associated with IFNα, while attenuating the TGFβ-associated TP63 response. Surprisingly, STAT3 activity was inhibited under IFNα, but was higher than under DMSO. This may be the result of studying the cells under a high fat media condition, generally resembling more diseased cells (which can be seen generally inflamed or “fibrotic” in Fig. 7b-C).

At the phenotype level, IFNα-treated spheroids had lower MAS and fibrosis-associated scores than the DMSO condition, whereas TGFβ-treated spheroids had higher scores (Fig. 7B-C). Both the gene expression and TF activity scoring placed the combined condition between the two single-cytokine conditions. In the gene expression-inferred phenotypes, scores were close to DMSO, while the separation of the single-cytokine conditions was more pronounced in the TF activity-inferred scores. These directional relationships are consistent with the opposing disease-associated transcriptional shifts predicted by the receptor-stimulation simulations. They do not constitute direct measurements of steatosis, inflammation, or fibrosis in the spheroids, as phenotypes were scored using the clinically trained PLSR model.

Altogether, the RNA profiles provide transcriptional support for selected model-derived responses, including TGFβ-associated TP63 activity and opposing IFNα- and TGFβ-associated phenotype scores.

## 3. Discussion

Here, we constructed an interpretable, genome-scale model of signaling trained on drug perturbations in primary human hepatocytes. We used it to ask how TGFβ- and IFNα-induced intracellular signaling interact in the context of MASH. Evidently from our results, the two receptor programs push the predicted MAS and fibrosis stage scores in opposite directions. When TGFBR1 and IFNAR1 were perturbed together, the response was competitive; i.e., the activation of one pathway offset the phenotypic effect of the other. Developing a gradient-based approach for the rational decomposition of the MoA of a perturbation into primary pathways allowed us to interrogate these responses. This pointed to a recurrent signaling module built around SMAD3, TP63, STAT2, and STAT3, and targeted perturbations within that module separated TFs that set the overall disease-associated state from those required to couple receptor activity to the competition itself. Taken together, the simulations describe one mechanistic route by which TGFβ and IFNα may jointly regulate disease-associated transcriptional states in human hepatocytes.

Before interpreting any of these results, however, we had to establish that the trained signaling networks carried transferable information and were not simply fitting the drug perturbation dataset. The DT-LEMBAS ensembles predicted the activity of 99 out of the 104 TFs more accurately than models trained with shuffled outputs, and retained predictive capacity when individual drug doses were withheld from training. The Leave-One-Drug-Out (LODO) validation was the more stringent test for our purposes, since it interrogates the signaling component that the subsequent simulations actually use. With an entire drug removed from training, stimulation of its annotated targets partially reproduced its measured TF response. Recovery of canonical responses following TGFBR1/2 and IFNAR1/2 perturbation added an in silico biological validation before any experiment. Comparing against a linear dose response model, DT-LEMBAS was not uniformly superior across every TF, time point, and dose; however, this may be due to this kind of response data being able to be explained by linear effects, rather than the model failing to capture useful information. Thus, the linear comparison can only indicate for which TFs, indeed, a non-linear architecture is required to increase performance in predictions. Therefore, our computational model is able to extrapolate to unseen doses and even in silico perturbations, in a statistically significant way. This network-constrained model, thus, offers a biologically structured, non-linear signaling system that can be perturbed and decomposed to generate mechanistic hypotheses.

With the models validated at this level, TGFBR1 and IFNAR1 produced opposing changes in the predicted clinical phenotypes. Loss of TGFBR1 reduced both MAS and fibrosis stage scores, whereas loss of IFNAR1 or IFNAR2 increased them. The direction of these shifts is consistent with the established contribution of TGFβ signaling to fibrotic and metabolic liver pathology, while it suggests that IFNα-responsive signaling can counterbalance this disease-associated program (Dooley & ten Dijke, 2012; Inagaki *et al*, 2003; Wieser *et al*, 2018). The two-dimensional stimulation analysis showed that neither receptor accounts for the behavior on its own. Perturbations of opposite direction reinforced each other, whereas co-directional perturbations partially cancelled and remained closer to the untreated state. What sets the phenotype-associated transcriptional state in these simulations is the balance between the two receptor programs.

These findings extend earlier observations of antagonism between interferon- and TGFβ-mediated signaling. Competition has been demonstrated for IFNγ in reduced experimental systems (Ulloa *et al*, 1999), with supporting observations in preclinical liver models (Weng *et al*, 2007; Park *et al*, 2025), and related interactions having been reported in other cellular and disease contexts (Bright & Sriram, 1998; Biffi *et al*, 2019). More recently, we showed that transcriptomic and phenotypic experiments indicated direct antagonism between interferon- and TGFβ-associated programs in human liver 3D models (Cadavid *et al*, 2026). We complement these studies by placing the interaction within a genome-scale signaling network trained on primary human hepatocyte responses. One caveat worth stating is that the present simulations demonstrate functional competition between predicted receptor programs, and not direct biochemical inhibition between the receptors or their immediate effectors. The sequential simulations further suggest that this balance may depend on the order in which the signaling states are established. That observation remains a model-derived hypothesis, but it raises the possibility that prior exposure to one cytokine alters how the hepatocyte responds to the other.

Decomposition of the simulated responses returned SMAD3, TP63, STAT2 and STAT3 as recurrent components of the TGFBR1–IFNAR1 interaction at 2, 8 and 24 h. Experimental validation on in vitro liver models confirmed the importance of these signaling nodes. SMAD3 and STAT2 provide canonical links to TGFβ and interferon signaling, respectively, while STAT3 has broader and context-dependent functions in metabolic and inflammatory liver disease (Liu *et al*, 2023). The mediator-knockout simulations helped separate these roles. SMAD3 loss strongly reduced the absolute predicted MAS and fibrosis stage scores, consistent with its established contribution to TGFβ-driven liver pathology and systemic metabolic regulation (Tan *et al*, 2011; Xu *et al*, 2016; Deng *et al*, 2024). STAT2 loss instead removed the protective effect associated with IFNα signaling. TP63 loss attenuated the disease-promoting response to TGFβ and narrowed the separation between opposing receptor states, most clearly for MAS. STAT3 and TP53 perturbations mainly shifted the overall phenotype level without collapsing the competitive response to the same degree. This scan therefore distinguishes regulators of the absolute disease-associated state from mediators that transmit the opposing effects of the two receptors. Within that distinction, STAT2 and TP63 emerge as complementary candidate mediators of the competitive interaction, while SMAD3 remains a major determinant of the TGFβ-associated phenotype.

TP63 was the least expected member of this core module. The p63 transcription factor has multiple isoforms with distinct and sometimes opposing functions, and TAp63 has previously been implicated in the control of hepatic steatosis and metabolic homeostasis in mouse models (Porteiro *et al*, 2017; Fondevila *et al*, 2024). However, its contribution to TGFβ-responsive signaling in human hepatocytes, however, has not been established. Additionally, the TP63 gene was not overexpressed in the hepatocyte data examined here. This is an important qualification because the inferred TF activity may arise from overexpression or just activity (i.e., phosphorylation of some isoform of p63). Thus, the latter is a more plausible explanation, as TP53 TF activity is indeed induced (according to the inference) in the examined data, while evidence obtained from the supplementary single-cell analysis also supports its relevance despite the low expression in hepatocytes. Its recurrence in the decomposed paths, together with the selective effect of its simulated knockout, suggests that it may connect TGFβ signaling to the metabolic component of the predicted disease state. We keep this interpretation tentative, because neither the TF activity inference nor the current network representation distinguishes TP63 isoforms. Isoform-resolved measurements will be needed to determine whether a specific p63 program mediates the interaction and whether its role is conserved in human liver tissue.

Methodologically, this work addresses a persistent problem with genome-scale neural signaling models: they can represent complex non-linear and recurrent interactions, but their internal mechanisms resist the interpretation approaches designed for linear or acyclic networks. Our Jacobian and chain-rule decomposition evaluates the local influence of individual edges under defined receptor-stimulation states, which allowed state-dependent paths to be recovered from the full network. Aggregating paths across independently trained models and across time points prioritized recurrent mechanisms and limited our reliance on a single fitted parameterization. The resulting compact networks reproduced most of the directional phenotype response of the complete models, though not all of it, and are better read as explanatory reductions than as exact substitutes for the original networks. Convergence with an orthogonal decomposition based on the Gaussian Graph Modeling approach (Dobra *et al*, 2004; Pihur *et al*, 2008) supports the same four factors. While convergence is not causality, it makes it less likely that the selection is an artifact of one path-scoring assumption.

Connecting the simulated TF states to PLSR models derived from human clinical data adds a translational layer. This projection allowed the consequences of receptor perturbations to be expressed as transcriptional variation associated with MAS and fibrosis stage, instead of relying only on canonical pathway markers. These scores are not measurements of steatosis or fibrosis. They are disease-associated transcriptional states inferred from TF activity, and their accuracy depends both on the clinical model and on how well drug-trained primary hepatocyte networks stand in for diseased liver tissue. Even so, the framework offers one practical way to join mechanistic signaling models to human disease data, and it may narrow the gap between what reduced experimental systems show and what is observed in patients (Teufel *et al*, 2016; Santhekadur *et al*, 2018; Vacca *et al*, 2024).

The competitive organization of the two pathways also carries a therapeutic implication. Inhibiting TGFβ signaling is an attractive strategy for reducing fibrotic progression, but systemic interference with such a pleiotropic pathway can be costly in adverse effects. Our results suggest that the phenotype-associated output of TGFβ signaling might also be modified through the opposing IFNα–STAT2 program, or through downstream mediators such as TP63. The converse follows as well: suppressing interferon-responsive signaling could inadvertently favour a more disease-associated hepatocyte state. None of this establishes the drugs used for model validation as therapeutic candidates. Compounds associated with elevated predicted TGFBR1/2 or IFNAR1/2 activity were used to examine whether the model recovered the expected signaling states, and not as evidence of receptor binding, clinical efficacy, or repurposing potential. The nodes we identify will have to be evaluated experimentally before their therapeutic relevance can be judged. Finally, this study provides a proof-of-concept on how these genome-scale neural networks representing intracellular signaling can be used to explore many in silico perturbations, and thus more candidate signaling nodes could be identified in the future to modulate the advancement of the disease.

The study’s main limitation lies in the data used to train the model using only primary human hepatocytes. The models describe hepatocyte-intrinsic signaling, whereas MASH develops through interactions among hepatocytes, hepatic stellate cells, immune populations, and endothelial cells. A hepatocyte model cannot reproduce the multi-cellular production, clearance, and feedback regulation of TGFβ and interferons in diseased liver tissue. Moreover, the receptor inputs are expressed on an arbitrary simulation scale and should not be read as biological cytokine concentrations, and the separately trained 2, 8 and 24 h ensembles describe time-specific response states rather than a continuous dynamical trajectory. Both the TF activities and the clinical phenotypes are inferred/calculated quantities, so errors or biases in the regulon, the prior-knowledge network, drug-target annotations, and the clinical PLSR model propagate into the simulations. The clinical PLSR model itself is trained on activity inferred by bulk RNA-seq of liver samples. This means that maybe the observed activity is coming from measured transcripts of other cell types too, even though it is safe to assume the observed transcripts are mainly coming from hepatocytes. Finally, the reduced-path models did not reproduce every response of the complete ensembles, and the MoA decomposition adds a further interpretational limitation. Its Jacobian describes local behavior around the simulated state, while path pruning and the incomplete prior network affect which mechanisms are retained. The selected paths should therefore be considered recurrent model-derived hypotheses, rather than the unique or definitive biological routes.

Overall, our results describe TGFβ and IFNα as competing genome-scale signaling programs rather than independent inputs to the hepatocyte disease state. Interpretable decomposition of the DT-LEMBAS ensembles connected this competition to a compact and experimentally testable module involving SMAD3, TP63, STAT2 and STAT3. Beyond the specific interaction studied here, the approach provides a general strategy for extracting mechanistic hypotheses from nonlinear, genome-scale signaling models and linking them to clinically associated transcriptional phenotypes. The framework can serve as a decision layer between perturbational data and experimental design. By screening perturbations and reducing selected responses to a tractable set of candidate mediators, it can focus resource-intensive studies on a biologically motivated search space without treating inferred paths as definitive mechanisms.

## 4. Methods

### 4.1. Data and preprocessing

#### 4.1.1. Gene expression retrieval and preprocessing

Microarray gene expression data, containing 120 unique drugs each tested on hepatocytes for three doses and three time points, were retrieved from the Japanese toxicogenomics project (Uehara *et al*, 2010; Igarashi *et al*, 2015). The raw microarray gene expression data were normalized using the Robust Multichip Averaging (RMA) algorithm (Irizarry *et al*, 2003) included in the *affy* R package (Gautier *et al*, 2004). The aggregation of probes into genes is performed by taking the median expression across probes. Subsequently, the gene expression data are standardized by subtracting the mean per gene.

#### 4.1.2. Transcription Factor activity inference

Transcription factor (TF) activity is inferred using the VIPER algorithm (Alvarez *et al*, 2016a) coupled with the Dorothea regulon (Garcia-Alonso *et al*, 2019). TFs with high variance across technical replicates were filtered to ensure we kept only high-quality estimations of TF activity, and then we filtered technical replicates that were not correlated enough with the other replicate signatures, using an approach described in detail in previous work (Meimetis *et al*, 2025). Briefly, we first build a permutation-based null distribution of TF activity variance by calculating the variance of the activity of each TF across groups of pseudo-replicates. A one-tailed Kolmogorov-Smirnov statistical test is used to estimate whether the true variance is less than random, and thus filter TFs when this is not statistically true. To filter replicates, we build a null distribution of random correlations between TF activity profiles, using random sampling. We conduct this for every possible number of replicates within a sample, and then count how many random correlations are equal to or higher than the mean correlation of the technical replicates. Finally, replicates are merged by taking the median of the activity of a TF across replicates.

#### 4.1.3. Prior knowledge (PKN) signaling and Drug-Target Interaction (DTI) network construction

Drugs’ SMILES and known DTIs were retrieved from the pre-processed data of the Broad Institute’s Repurposing Hub (Corsello *et al*, 2017) and DrugBank (Wishart *et al*, 2006) used in the original DT-LEMBAS publication (Meimetis *et al*, 2024), as well as the ChEMBL database (Gaulton *et al*, 2017). For drugs where no DTIs were retrieved automatically, interactions were manually curated from PubChem (Kim *et al*, 2016) and DrugBank’s web interface. The published DT-LEMBAS protocol (Meimetis *et al*, 2025) was ultimately followed to pre-calculate the chemical similarity between drugs and construct the final prior knowledge network of intracellular interactions and DTIs that served as the scaffold for training, with the log-scaled dose used as input to train models (*dose_scale_* = log_10_(*dose* + 1)).

#### 4.1.4 Retrieval and preprocessing of clinical datasets of Metabolic Dysfunction-Associated Steatohepatitis (MASH)

RNA-seq. data of liver samples from human patients, across various stages of MASH and fibrosis, were retrieved from the Govaere et al. publicly available transcriptomic dataset (Govaere *et al*, 2020). Gene count data were downloaded from the ARCHS4 database (Lachmann *et al*, 2018). Counts per million (CPM) normalization and subsequent log2 transformation were applied. The gene expression matrix was centered (by subtracting the mean per gene), and then TF activity was inferred as already described in section 1.2.

### 4.2. Interactome-based deep learning modeling of hepatocyte signaling

#### 4.2.1. Model architecture and training

A GPU implementation of the DT-LEMBAS architecture (using the GPU version of the LEMBAS signaling network (Nordenstorm *et al*, 2026)) was trained for 1000 epochs to predict the activity of 104 TFs given the concentration of a drug, within a pre-defined drug-target space of 120 drugs and 134 potential targets. Because of the small DTI space, inference of new targets was not regularized at all. A batch size of 35 was used together with a momentum of 0.2 in the batch normalization layer of the drug module and a dropout rate of 0.1. The remaining parameters were left at their default values (as described in the published protocol (Meimetis *et al*, 2025)), except for the maximum number of recurrent neural network iterations, set to 90, and the tolerance of the spectral loss, set to 10⁻⁵. After validating the performance of a single model, an ensemble of 50 independently initialized models was trained for use in all downstream simulation and decomposition analyses, allowing model uncertainty to be propagated through later results. One ensemble of 50 models was trained for each of the three timepoints (resulting in three ensembles of 50 models).

#### 4.2.2. Model validation

The model was evaluated across three complementary validation schemes: i) classic 10-fold Cross-Validation (CV), ii) Leave-One-Out Cross-Validation (LOOCV), by leaving a whole dose level (low, medium, or high) out of the training set, iii) Leave-One-Drug-Out validation (LODO). Validation was performed separately for each time point. Moreover, it is worth noting that the exact performance of DT-LEMBAS is not of great interest, as it is simply required to know that the models’ weights are generalizable enough (even if that generalization is linear at the end) to be trusted for downstream analyses utilized the signaling part of the model. This is why the 3^rd^ validation regime (even though challenging and slightly out of the scope of this work) is very important to estimate the ability of the model to extrapolate to unseen in silico perturbations.

##### 4.2.2.1. 10-fold Cross-Validation

We split our dataset using 10% of the data for testing and 90% for training, 10 times (10-fold), for each of the three time points separately. The only modification from classic 10-fold CV is that the CV algorithm is forced to keep each drug with at least one of its doses in the training set. Thus, this is considered the easiest validation regime, and we compare the performance of the hepatocyte-specific DT-LEMBAS models against a random baseline of re-training them with data where the TFs’ labels are randomly shuffled (shuffled Y) in the training set, while kept the same in the test sets.

##### 4.2.2.2. Leave One Dose Out Cross-Validation

To make sure the model can extrapolate to unseen doses of a drug (or interpolate to intermediate levels), we utilize an approach where, for all drugs, the low, medium, or the high dose is completely left out of training. Then we re-train a shuffled Y version to compare against a random baseline, but we additionally compare against linear interpolation or extrapolation for each drug for each TF. The latter is simply complementary to test whether our non-linear model explains the observed TF activity better than a linear baseline.

##### 4.2.2.3. Leave-One-Drug-Out validation (LODO)

To evaluate the ability of the model to extrapolate to completely unseen perturbations (for later in silico simulations), we devise a new validation scheme where we completely leave every drug out of the training once, and then artificially stimulate its targets in silico according to a mechanism inferred from the previously trained ensemble (and known mechanism of action). Using this in silico stimulation, the predicted transcription factor (TF) activity was compared to the measured drug-induced TF activity to test whether the trained signaling network could retrieve it. The whole validation approach consists of seven steps: 1) identification of drugs safe to be removed, 2) inference of the drug-target interaction sign, 3) ranking the targets based on their importance, 4) model re-training without a specific drug, 5) artificial stimulation of held-out drugs’ targets, and finally 6) selection of the strength of stimulation and dose matching. The algorithm for in silico drug stimulation by knowing some of their known targets (with or without knowledge of their binding affinity) could be its own study, and it is currently out of the scope of this work, especially when considering the unknown off-target effects of drugs.

Agreement between predicted and measured TF activity was quantified by Pearson correlation at three levels: i) per TF across drugs, ii) per drug across all TFs, and iii) overall across all drug-TF pairs. The overall correlation was used both to select the stimulation strength and as the summary metric reported in the main text.

For Step 1, removing certain drugs can disconnect part of the signaling network, leaving some TFs unreachable from any remaining input and precluding a valid comparison. We therefore restricted LODO to drugs that can be removed without such isolation. A drug was defined as safe to remove if, after its removal, every TF remained reachable in the prior-knowledge network (PKN) graph from the targets of at least one other drug. In the dataset used, 113 of the 120 drugs (94.2%) were safe to remove and thus used for validation.

For Step 2, applying a directional in silico stimulation requires knowing whether a drug activates or inhibits each of its targets. We inferred this from the pre-trained ensemble models rather than assuming it. For the linear drug layer of one DT-LEMBAS, the effective interaction weight (*w_d,t_*) of a drug (*d*) on a target (*t*) was calculated (using Equation 1) from the learned drug-target weight matrix (*A*), the learned drug-drug similarity matrix (*W_sim_*), and the chemical similarity scaling parameters (*S*), as they are used in the drug layer of DT-LEMBAS (Meimetis *et al*, 2024, 2025).

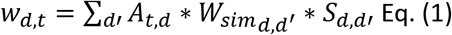

For an ensemble of 50 models, the consensus sign of interaction (d, t) was taken as the median of the signs of the weights calculated in Eq. (1). A positive consensus sign denotes activation (+1), a negative sign denotes inhibition (−1), and zero denotes an undetermined interaction. Targets with an undetermined sign received no stimulation.

For Step 3, to weight each target by how strongly a drug engages it (rather than stimulating all targets equally), we extracted a drug x target interaction score matrix for each model in the ensemble at each time point, using the established protocol for the linear drug layer to calculate the importance of a drug in affecting a target (Meimetis *et al*, 2025). Scores were then aggregated by their median across the ensemble and across the three timepoints, and then converted, per drug, to importance ranks on a 0–1 scale (percentile ranking). By default, similarity-propagated (“off-target”) interactions were excluded so that ranks reflected on-target engagement. These consensus ranks were used to scale the subsequent stimulation magnitude. It is worth noting that other rank normalizations were also tested (softmax and max normalization).

For Step 4, a single DT-LEMBAS model (rather than a full ensemble, for computational efficiency as there are 113 repeats per timepoint) was then re-trained per held-out drug using the same configuration as the main models. For each safe held-out drug *d*, the drug was removed from the drug-target matrix, the corresponding row and column were removed from the chemical-similarity matrix, and training samples in which d was the only active drug were excluded. The PKN structure was left unchanged.

For Step 5, the held-out drug’s effect was reproduced by stimulating all of its targets simultaneously. The input assigned to target *t* combined the inferred sign, a base stimulation strength *s*, and was scaled by the target’s consensus importance rank:

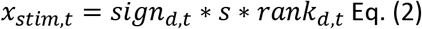

That way, the stimulation applied to the input node of target t is proportional to its importance rank (acting similarly to a binding affinity) and directed by its inferred sign. Targets with an undetermined sign received no stimulation. Moreover, an alternative approach was tested too, where ranks were not used, and all targets received the uniform stimulation value s (multiplied by their sign of interaction). Different stimulation strengths were tested, but the stimulation magnitude in general was kept moderate to avoid a knockout- or overexpression-like regime. TF activity was then read out at the network steady state, computed as for the main model.

Finally, for Step 6, because the artificial stimulation is dose-agnostic, each trained model was re-scored (without retraining) over a grid of base stimulation strengths s (as already mentioned in Step 5). Predicted TF activities were compared to the measured profiles, with negative predicted activities clamped to zero. As drug effects vary with concentration, for every drug the prediction was compared against the single experimental dose whose measured TF profile it best matched, selected either as the dose minimizing the mean absolute difference between predicted and true values, or, equivalently, as the dose maximizing the per-drug Pearson correlation across TFs. The base stimulation strength retained for reporting was the one maximizing the overall Pearson correlation between predicted and measured TF activity across all held-out drugs. This choice was verified to be consistent when strengths were ranked at the per-TF and per-drug levels.

### 4.3. In silico simulation and phenotype scoring

To inhibit the activity of a signaling node (or vice versa, activate the node) the trained signaling part of DT-LEMBAS is taken, and a negative value is assigned as input signal to the node. Then it propagates through the signaling network to ultimately predict downstream transcription factor activity. After investigating convergence of induced activities in a grid of in silico stimulations, knockdowns/knockouts are approximated as extreme inhibitions by assigning a numerical value of -3000. To generate a combinatorial grid of input stimulations for TGFBR1 and IFNAR1, we first generate a pool of 201 potential stimulation values ranging from -3000 (extreme inhibition) to 3000 (extreme activation), and we combine the two in panel of all the possible combinations (40401). These will be passed to predict induced TF activity. The TF activity is scored in terms of phenotypes’ scores (MAS and fibrosis stage) using a Partial Least Squares Regression (PLSR) model. Specifically, we trained a PLSR model on the calculated TF activity of clinical samples of different stages of the diseases, from a publicly available bulk RNA-seq datasets of human patients with MAFLD (Govaere *et al*, 2020). The TF activity was inferred as described in methods’ section 1.2. Ridge regression was only used when creating a network connecting disease nodes with TF nodes during MoA extraction for simplicity, but as it is show in prior literature there is not a significant difference in performance or predictions (Cadavid *et al*, 2026).

### 4.4. Network decomposition

To extract the subnetwork explaining (at most) the effect of IFNAR1 and TGFBR1 on MAS and fibrosis we utilize a pruning approach which consists of 6 steps. Note that, except for Step 1, these are repeated for each one of the 50 trained models in the ensemble.

In Step 1, we train a ridge regression model (using 10-fold CV, repeated 10 times) to predict MAS and fibrosis separately from the activity of transcription factors in the network, using the transcriptomic clinical dataset provided by Govaere et al(Govaere *et al*, 2020). We use the weights learned by the regression model to connect MAS and fibrosis to the intracellular signaling network. The transcription factor activity was calculated from gene expression using the using the VIPER method(Alvarez *et al*, 2016b) from the decoupleR resource(Badia-i-Mompel *et al*, 2022).

Ideally we aim to enumerate all possible paths from the receptors of interest to MAS and fibrosis, but since this is practically infeasible we perform an initial filtering of the edges in the network. Specifically, in Step 2, we build a 21×21 grid on artificial stimulation of IFNAR1 and TGFBR1, and for combinations we predict, through the signaling model and the ridge regression model, scores for MAS and fibrosis (denoted as *Y^_k_* with k=1 for MAS and k=2 for fibrosis). Then we calculate as a cost function the range of MAS and fibrosis induced by this grid as *L_k_* = max{*Y_k_*} − min{*Y_k_*}. Using this cost function for each phenotype we calculate its gradient with respect to every edge in the network, which signifies the importance of an edge in affecting the dynamic range of the phenotypes. We drop edges with a gradient less than 3*10^-2^ and 8*10^-3^ for MAS and fibrosis respectively. This way we immediately reduce the paths to a traceable number for enumeration.

In Step 4, we enumerate all paths (with a maximum length of 8) from the receptors to the phenotypes, for each of the 50 models in the ensemble. Even with initial pruning, the paths are too many to enumerate sometimes, and thus we begin scoring them (as it will be described in detail in Step 5) to prioritize some of them. Specifically, we set a maximum 1200 paths for each batch of enumeration. If the 1^st^ batch is less than that we finish enumeration for this model and proceed to the next in the ensemble. If this number is exceeded we score all 1200 paths and keep only the top 250 paths. The rest of the paths and all their edges are removed. Then we proceed to the next batch and repeat the process. We stop after 10 attempts or if we indeed enumerated all possible paths. This phenomenon only happens for a few of the models in the ensemble, thus, because each time we keep the top 250 paths from each batch, and also we have evaluated up to 12000 paths, we reason that we miss only a small amount of information, which is anticipated to be corrected by the ensemble.

In Step 5, we finally score all enumerated paths per model, utilizing an integrated gradient approach. Specifically, we once again define a grid of stimulations, and for each stimuli we define the input to the model as a stimuli, starting from a zero baseline to the actual stimulation: *x_a_* = *x_baseline_* + *α* ∗ (*x_stimulion_* − *x_baseline_*), with x_baseline_ being zero in our case, and α being a ratio from 0 to 1, generated by the steps selected for the integrated gradient calculation. Then we calculate the single-step Jacobian (gradient for each edge, i.e. gradient of each target node with respect to a source node) at the equilibrium (final step of the recurrent neural network) of the model for a given input x_a_ as:

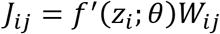

given that the activity of signaling node i is calculated as *z_i_* = *f*(*Wx*^∗^ + *bias*). x* is the activity of nodes at equilibrium, calculated by a full iteration pass of the model with input x_a_. f(x) is the activation function used by the neural network, W is the weight matrix of the signaling network and bias is the bias term as calculated in the LEMBAS and DT-LEMBAS models(Nilsson *et al*, 2022; Meimetis *et al*, 2024).

The Jacobian of the output scaling layer of LEMBAS is calculated using autograd (for a given TF we term this as J^TF^), while for the connection of MAS and fibrosis with TFs, this Jacobian are basically the weights of the ridge regression model (for a given TF we term this c^TF^).

The score for one step of the IG algorithm of whole paths is calculated using the chain rule, i.e. meaning for e.g. a path p = (IFNAR1 -> node1 -> node2 -> … -> TF -> MAS) the score is given by:

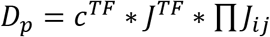

We accumulate across multiple integrated gradient steps to get the final path score:

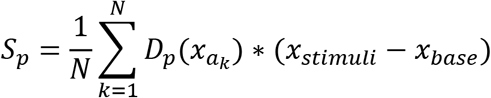

Because of the small numerical values of the Jacobian, we anticipate that larger paths (even though the maximum length is 8) can be penalized. For this purpose we utilize a geometric mean-based normalization along a path of length L_p_:

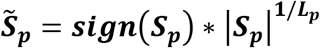

This last formula is equivalent to the geometric mean of the chain-ruled Jacobians along a path, and corrects for the length of a path.

Finally, in Step 6, we select the top 3 paths (based on *S̃^∼^_p_*) from each model of the ensemble and we assign a frequency score for each of the paths in the final aggregated network.

A complementary Gaussian Graphical Model (GGM) construction method is used to get more computational evidence on the validity of the extracted subnetworks. More details can be found in Appendix Supplementary Methods 2. Briefly, for each simulation, transcription factor and signaling node activity scores were used to estimate a precision matrix, and we utilized the Graphical Lasso (Glasso) algorithm implemented in Python.

### 4.5. Experimental validation (placeholders — fill as data arrive)

- Primary human hepatocyte spheroid culture and treatments
- Western blotting and pathway inhibitors
- RNA-sequencing and processing
- Phenotype imaging in co-culture
- Single-cell data analysis (Gribben et al.)

### 4.6. Statistics and reproducibility

Statistical analyses were performed in R (packages rstatix, stats) and Python (scipy.stats). Comparisons of model performance and of predicted activities between conditions were assessed with two-sided Wilcoxon rank-sum (Mann–Whitney) tests, except where a directional hypothesis was explicitly tested (e.g. suppression of marker-TF activity below control under in silico knockout), for which a one-sided Wilcoxon test was used. Whole distributions were compared with the non-parametric Kolmogorov-Smirnov (KS) test, including the one-tailed KS test used to filter high-variance TFs during preprocessing. Agreement between predicted and measured TF activity (dose-extrapolation and LODO validations) was quantified by Pearson’s correlation, with model performance benchmarked against label-shuffled (“shuffled-Y”) null models. Where multiple TFs were tested in parallel, p-values were adjusted for multiple comparisons using the Benjamini-Hochberg (BH) procedure. Unless otherwise noted, *n* denotes the ensemble of 50 independently trained models per timepoint. The unit of comparison (model, TF, drug, or replicate) is stated in each figure legend. Shaded bands denote ±1 standard error of the mean (SE). In all boxplots, the center line denotes the median, the box bounds the 1^st^ and 3^rd^ quartiles, and the whiskers points within 1.5× the interquartile range (IQR). Significance is denoted as ****p ≤ 10⁻⁴, ***p ≤ 10⁻³, **p ≤ 10⁻², *p ≤ 0.05, and ns for p > 0.05.

## 5. Data availability

The raw microarray data used to train our models can be found in ArrayExpress database hosted in EBI (accessed in 2025) under the E-MTAB-798 accession number. The preprocessed microarray data can be accessed and downloaded from the GitHub repository of the study: https://github.com/NickMeim/LiverLEMBAS. The clinical dataset (Govaere *et al*, 2020), used for training models to score transcriptional profiles in terms of clinical phenotypes, can be retrieved from the Gene Expression Omnibus (GEO) with the accession number GSE135251. Drug SMILES and known drug-target interactions were obtained from the Broad Institute Repurposing Hub, DrugBank, and ChEMBL, with additional interactions manually curated from PubChem and DrugBank. The assembled drug-target mask, SMILES, chemical-similarity matrices, and the prior-knowledge signaling network are available in the study’s GitHub repository (https://github.com/NickMeim/LiverLEMBAS). The subset of data used from the single-cell dataset (Gribben *et al*, 2024) is available in its study’s R Shiny app: https://www.mohorianulab.org/shiny/vallier/LiverPlasticity_GribbenGalanakis2024/. All the trained DT-LEMBAS models are deposited on Zenodo. All analyzed data that were used to train our models and produce all tables and figures are available at https://github.com/NickMeim/LiverLEMBAS as well. The code to generate and pre-process data, figures, and tables, as well as train models, is available in at the same GitHub repository.

## 6. Author contributions

N.M. and D.A.L. conceptualized and supervised the study. N.M. implemented the code and executed the simulations, preprocessed the data, trained the final models, designed the case studies and computational experiments, and analyzed their results. T.M. run the GMM analysis. T.M. and X.M. cultured cells and run perturbation experiments to measure RNA and activity of specific proteins, together with imaging organoids, under supervision of L.G.G. L.F. took care of RNA purification and sample preparation for RNA-seq, and helped with cell culturing. N.M. wrote the manuscript. N.M., T.M. and D.A.L edited the manuscript.

## 7. Disclosure and competing interest statement

The authors declare no competing or non-competing financial interests related to the publication of this study.

## 8. Acknowledgements

The authors would like to thank Brian Joughin, Hratch Baghdassarian, Luka Karginov, Christine Wiggins, Jose Cadavid, and Erin Tevonian for their valuable input on this work. We are grateful for funding support from NovoNordisk and the Army Institute of Collaborative Biotechnologies collaborative agreement W911NF-19-2-0026. NM would also like to acknowledge funding support from the MIT Takeda Fellowship program.

## Supporting information

Supplementary Matterial

## Notes

https://github.com/NickMeim/LiverLEMBAS

### Competing Interest Statement

The authors have declared no competing interest.

