## Supplementary Matterial for "Interactome-based deep learning and path-level decomposition reveal TGFβ-IFNα competition in MASH"

### Supplementary Figures

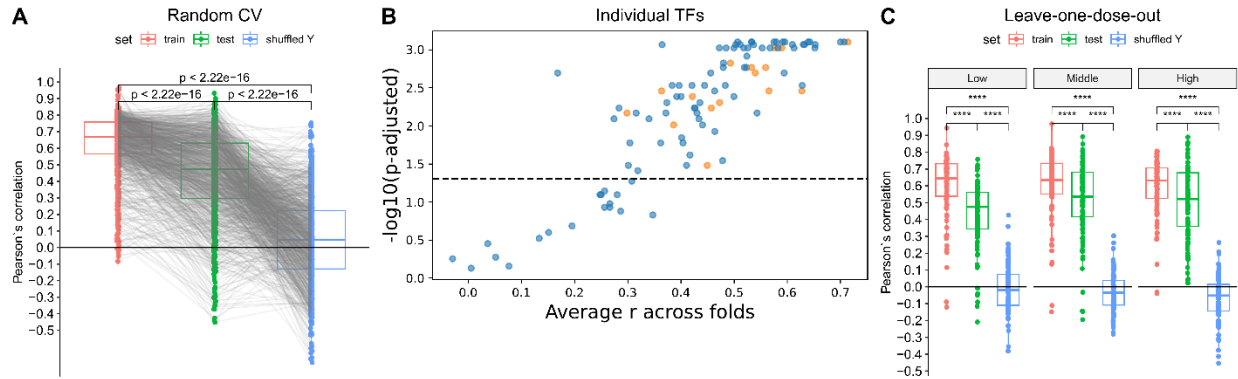

**Figure S1: Predictive performance of DT-LEMBAS at 2 hours.** **A)** Pearson correlations between predicted and observed transcription factor (TF) activities in random 10-fold cross-validation. Points represent TF-fold pairs. **B)** Scatterplot illustrating the mean test correlation per TF and BH-adjusted p-value from comparing it with a random (shuffled) baseline, using a one-sided Mann–Whitney U test. Orange points mark phenotype-associated TFs. **C)** Leave-one-dose-out performance for each TF, withholding low, middle, or high doses. Shuffled-Y models use randomized training TF labels. Comparisons in A and C use paired two-sided Wilcoxon tests. Boxes show medians and interquartile ranges. \*\*\*\*p-value $\leq 0.0001$.

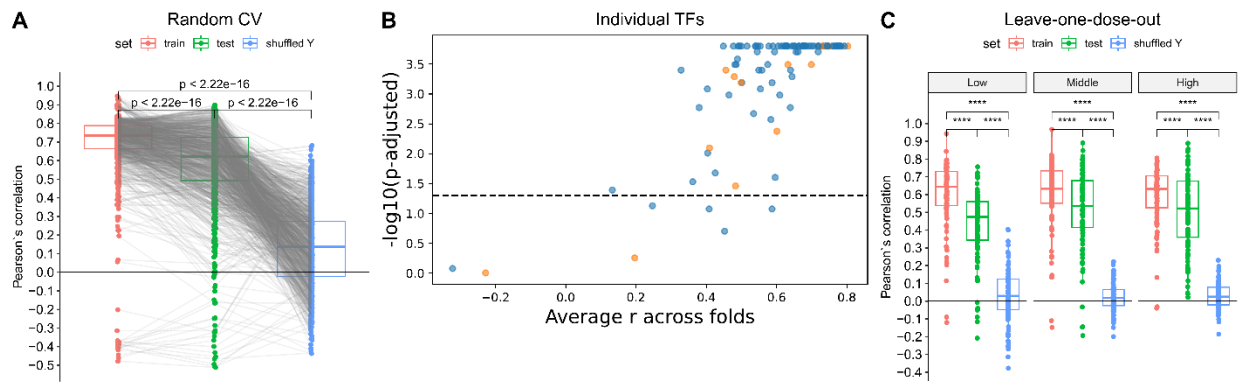

**Figure S2: Predictive performance of DT-LEMBAS at 24 hours.** **A)** Pearson correlations between predicted and observed transcription factor (TF) activities in random 10-fold cross-validation. Points represent TF-fold pairs. **B)** Scatterplot illustrating the mean test correlation per TF and BH-adjusted p-value from comparing it with a random (shuffled) baseline, using a one-sided Mann–Whitney U test. Orange points mark phenotype-associated TFs. **C)** Leave-one-dose-out performance for each TF, withholding low, middle, or high doses. Shuffled-Y models use randomized training TF labels. Comparisons in A and C use paired two-sided Wilcoxon tests. Boxes show medians and interquartile ranges. \*\*\*\*p-value $\leq 0.0001$.

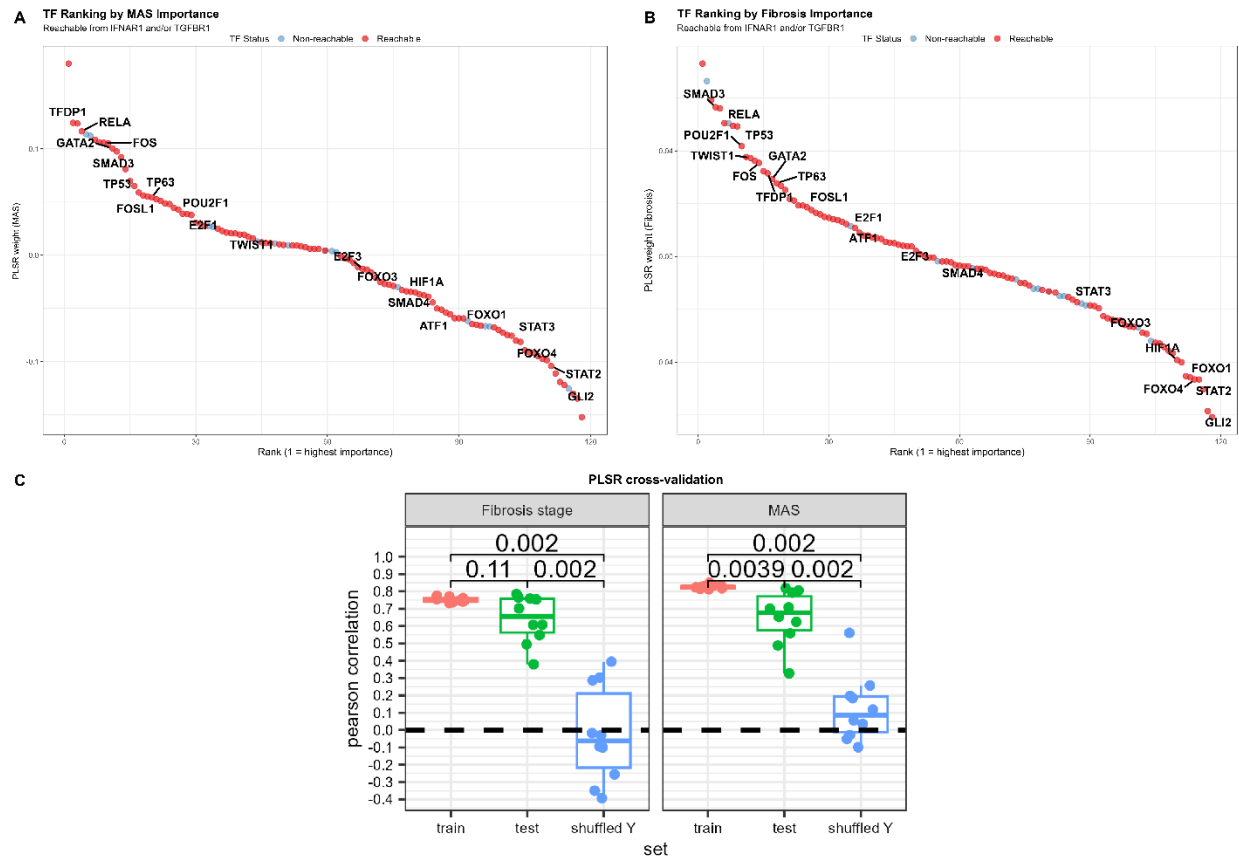

**Figure S3: Clinical phenotype projection and TF weights.** A, B) Signed partial least-squares regression (PLSR) coefficients fitted to all inferred human liver TF activities for MAS (A) and fibrosis stage (B), ranked from highest to lowest. Red TFs are reachable from IFNAR1 and/or TGFBR1, while blue TFs are not. C) 10-fold cross-validation of the clinical phenotype projection using the 104 DT-LEMBAS TFs. Training, test, and shuffled-Y Pearson correlations are compared using paired two-sided Wilcoxon tests. Boxes show medians and interquartile ranges.

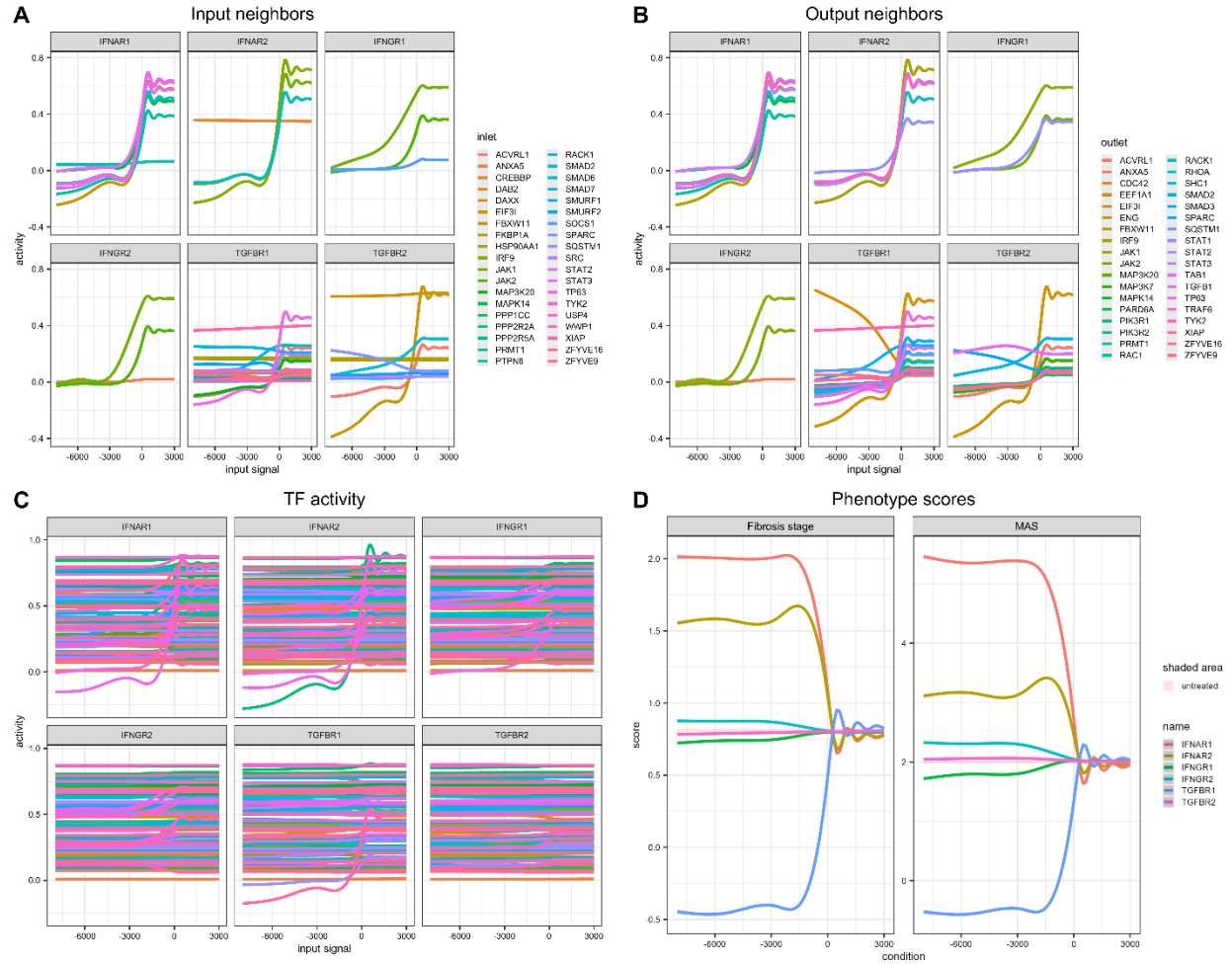

**Figure S4: Convergence of receptor perturbations at 2 hours.** **A, B)** Activities of incoming (**A**) and outgoing (**B**) neighboring nodes across receptor input strengths. **C)** Predicted TF activities across receptor input strengths. **D)** PLSR-predicted MAS and fibrosis stage scores across receptor input strengths. Curves summarize simulations across 50 models; ribbons show 95% confidence intervals of smoothed fits. Shading denotes the untreated mean $\pm 1$ SE. Negative and positive inputs represent inhibition and activation, respectively, in arbitrary units.

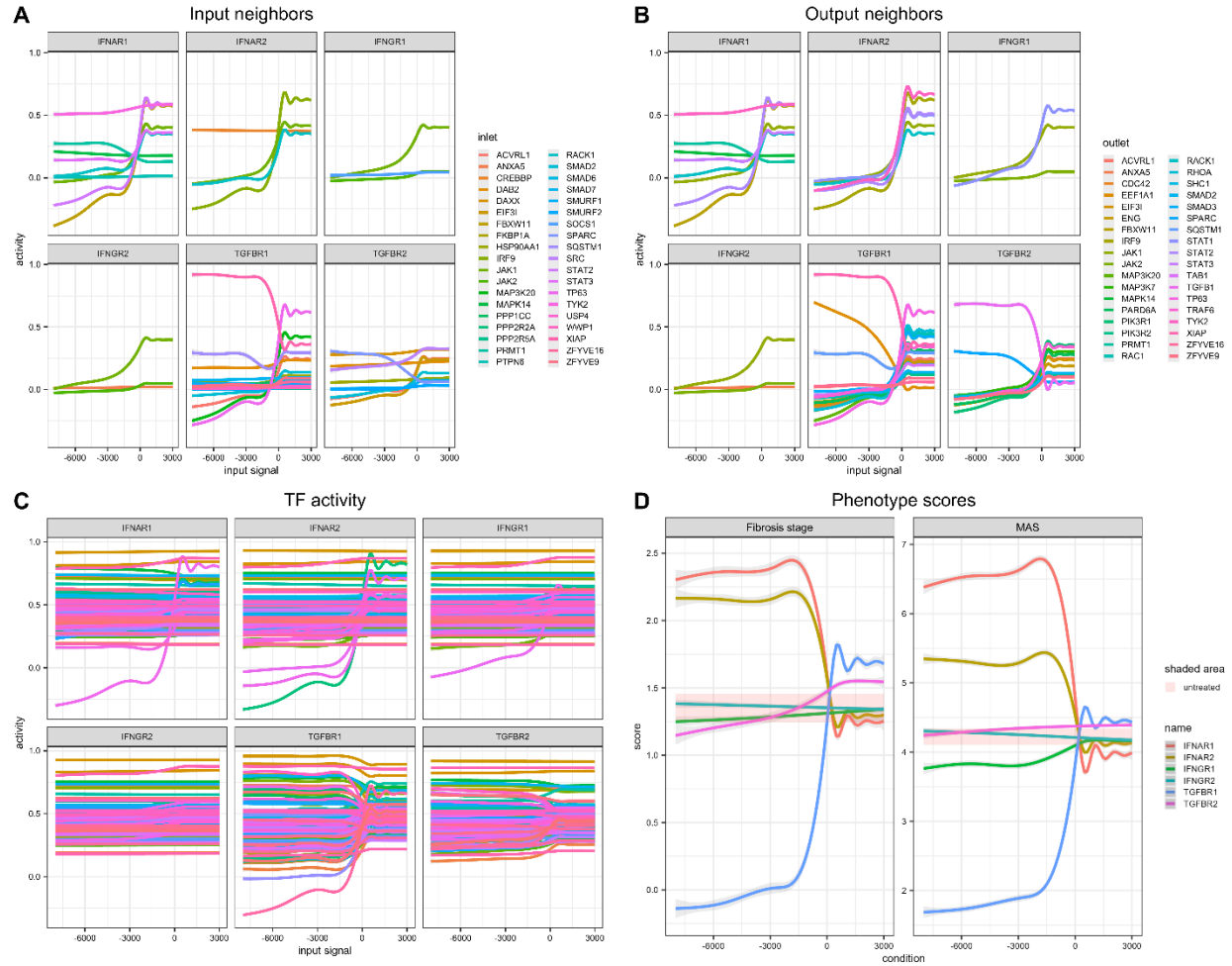

**Figure S5: Convergence of receptor perturbations at 8 hours.** **A, B)** Activities of incoming (A) and outgoing (B) neighboring nodes across receptor input strengths. **C)** Predicted TF activities across receptor input strengths. **D)** PLSR-predicted MAS and fibrosis stage scores across receptor input strengths. Curves summarize simulations across 50 models; ribbons show 95% confidence intervals of smoothed fits. Shading denotes the untreated mean $\pm 1$ SE. Negative and positive inputs represent inhibition and activation, respectively, in arbitrary units.

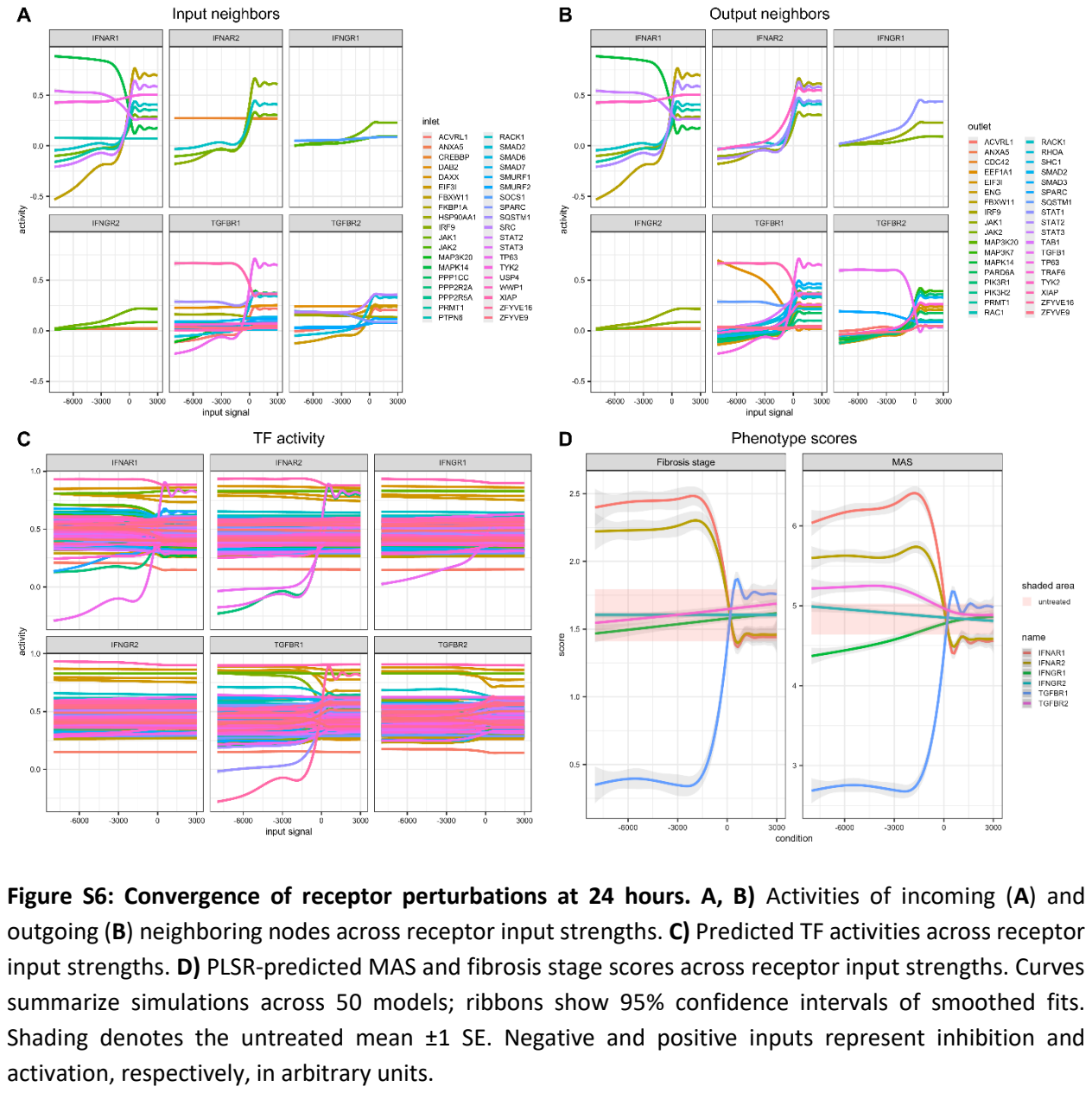

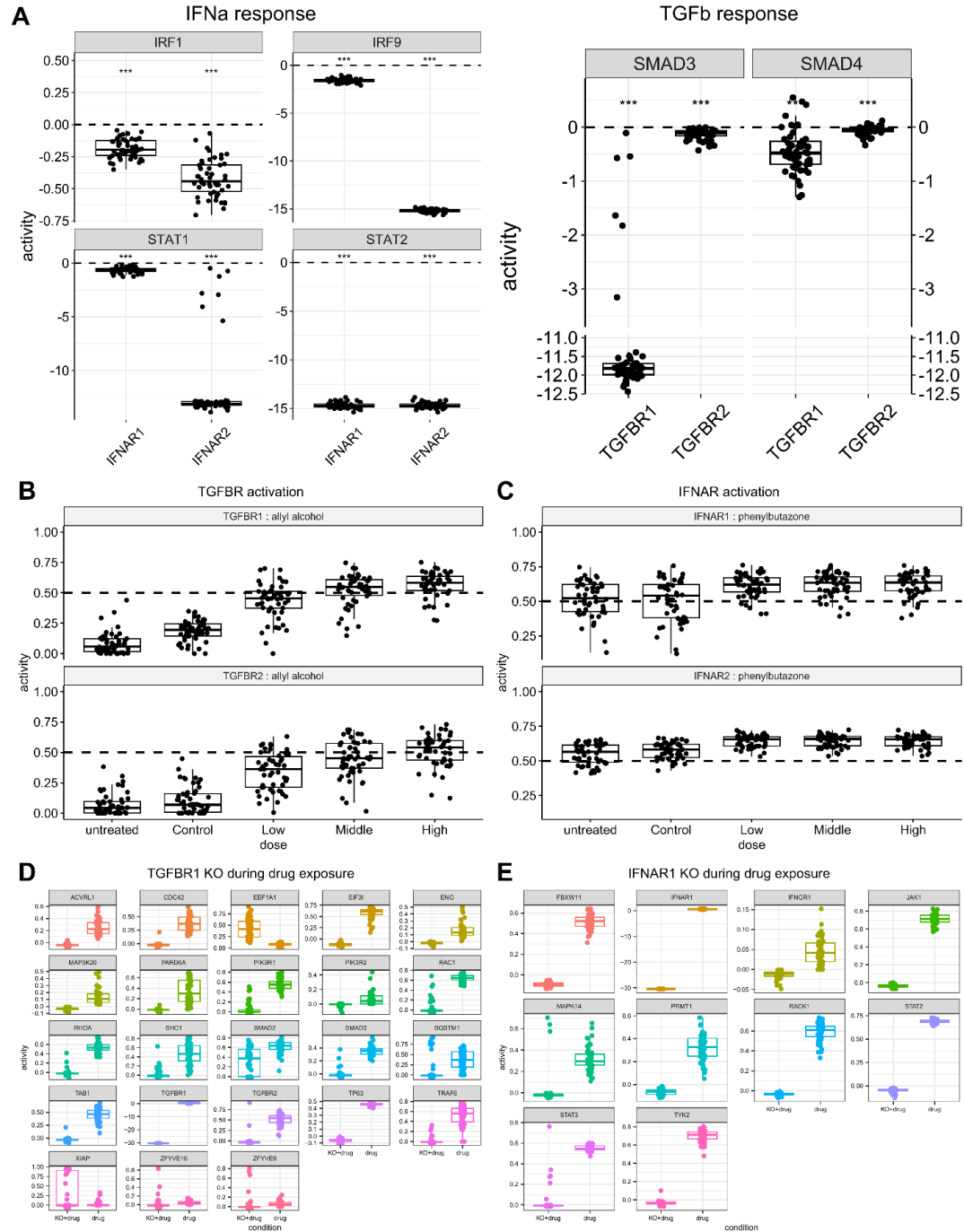

**Figure S7: Receptor knockout and drug-induced signaling at 2 hours. A)** Changes in IFNα-responsive and TGFβ-responsive TF activities after receptor knockout, relative to untreated models. One-sided one-sample Wilcoxon tests assess decreases from zero. **B, C)** TGFBR1/2 activities under allyl alcohol (**B**) and IFNAR1/2 activities under phenylbutazone (**C**), across the indicated controls and doses. These drugs were model-identified activators of the receptors. **D, E)** Neighboring nodes' activities during the corresponding drug exposures with or without TGFBR1 (**D**) or IFNAR1 (**E**) knockout. Points represent 50 independently trained models. Boxes show medians and interquartile ranges. \*\*\*p-value < 0.001; ns, p-value ≥ 0.05.

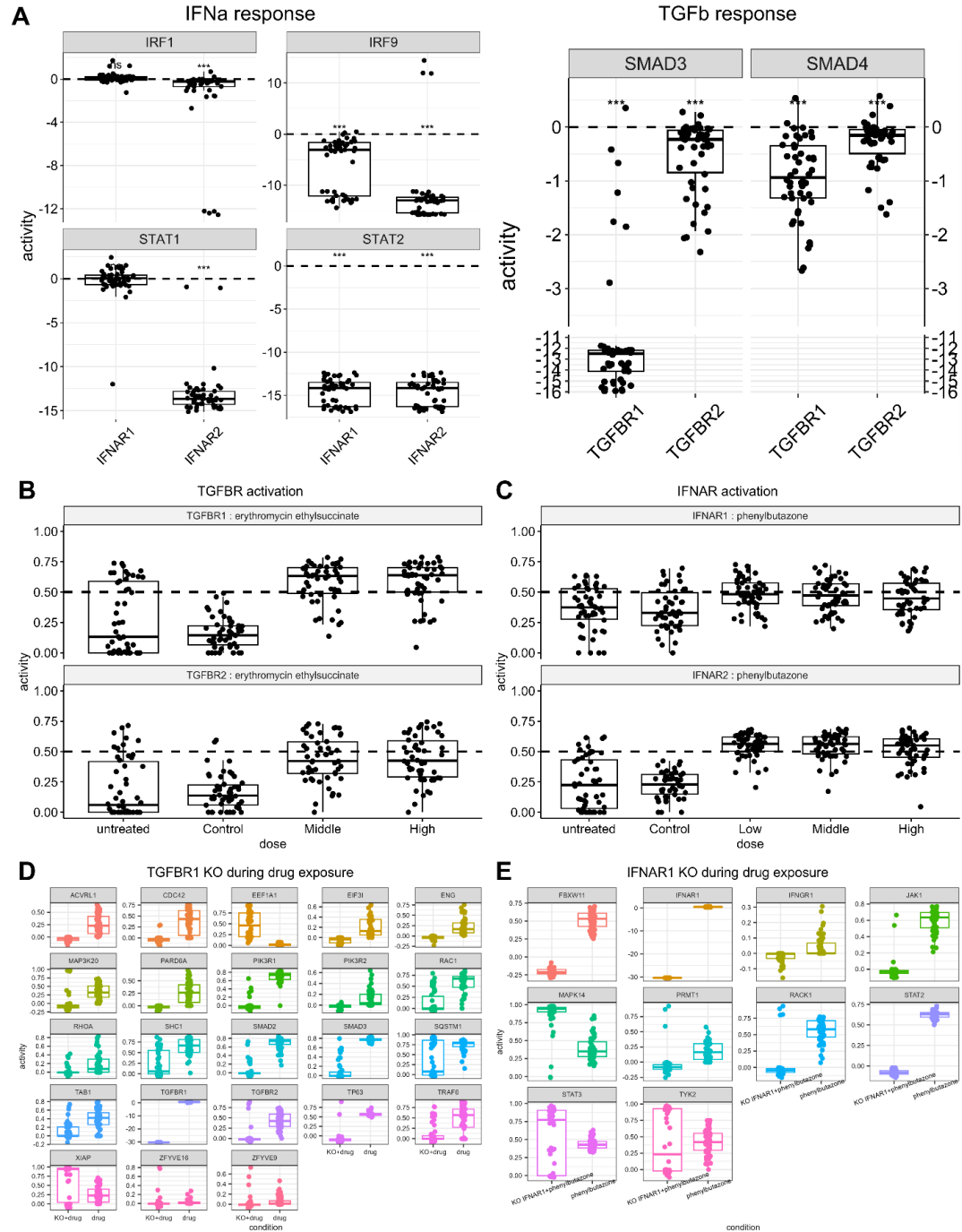

**Figure S8: Receptor knockout and drug-induced signaling at 24 hours.** **A)** Changes in IFNα-responsive and TGFβ-responsive TF activities after receptor knockout, relative to untreated models. One-sided one-sample Wilcoxon tests assess decreases from zero. **B, C)** TGFBR1/2 activities under erythromycin ethylsuccinate (**B**) and IFNAR1/2 activities under phenylbutazone (**C**), across the indicated controls and doses. These drugs were model-identified activators of the receptors. **D, E)** Neighboring nodes' activities during the corresponding drug exposures with or without TGFBR1 (**D**) or IFNAR1 (**E**) knockout. Points represent 50 independently trained models. Boxes show medians and interquartile ranges. \*\*\*p-value < 0.001; ns, p-value ≥ 0.05.

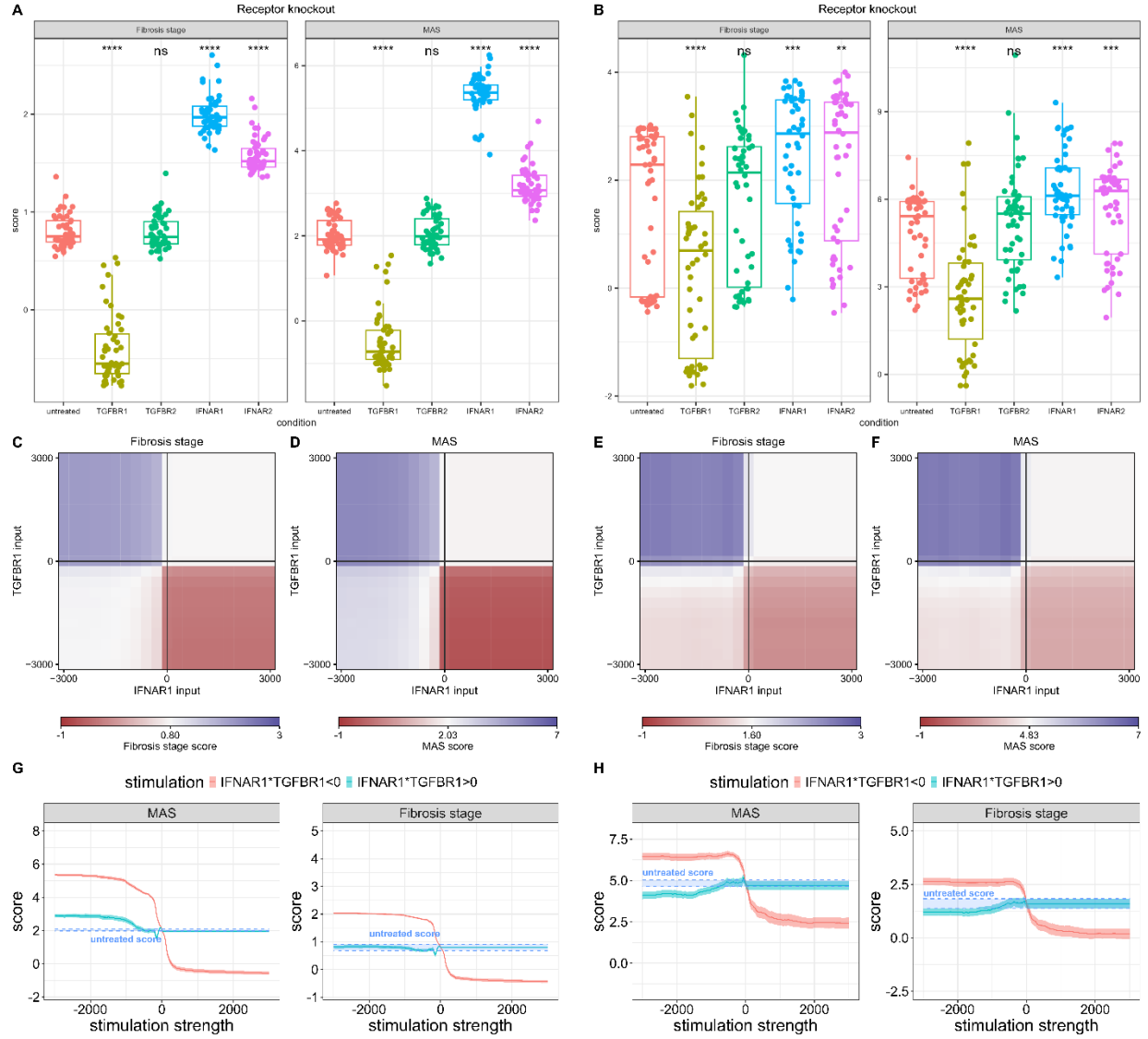

**Figure S9: Combined TGFBR1–IFNAR1 perturbations at 2 and 24 hours.** **A, B)** Receptor-knockout phenotype scores at 2 (**A**) and 24 (**B**) hours; points represent 50 models. Two-sided Wilcoxon rank-sum tests compare knockouts with untreated models. **C–F)** Ensemble-mean fibrosis stage and MAS scores across receptor input grids at 2 (**C, D**) and 24 (**E, F**) hours. Color scales are centered on untreated scores. **G, H)** Scores along opposing-sign (coral) and co-directional (turquoise) diagonals at 2 (**G**) and 24 (**H**) hours. Stimulation strength denotes IFNAR1 input. Lines show means and colored ribbons $\pm 1$ SE. Dashed lines and grey shading mark untreated references. Inputs are in arbitrary units. Boxes show medians and interquartile ranges. \*\*p-value < 0.01, \*\*\*p-value < 0.001, \*\*\*\* p-value $\leq 0.0001$; ns, p-value $\geq 0.05$.

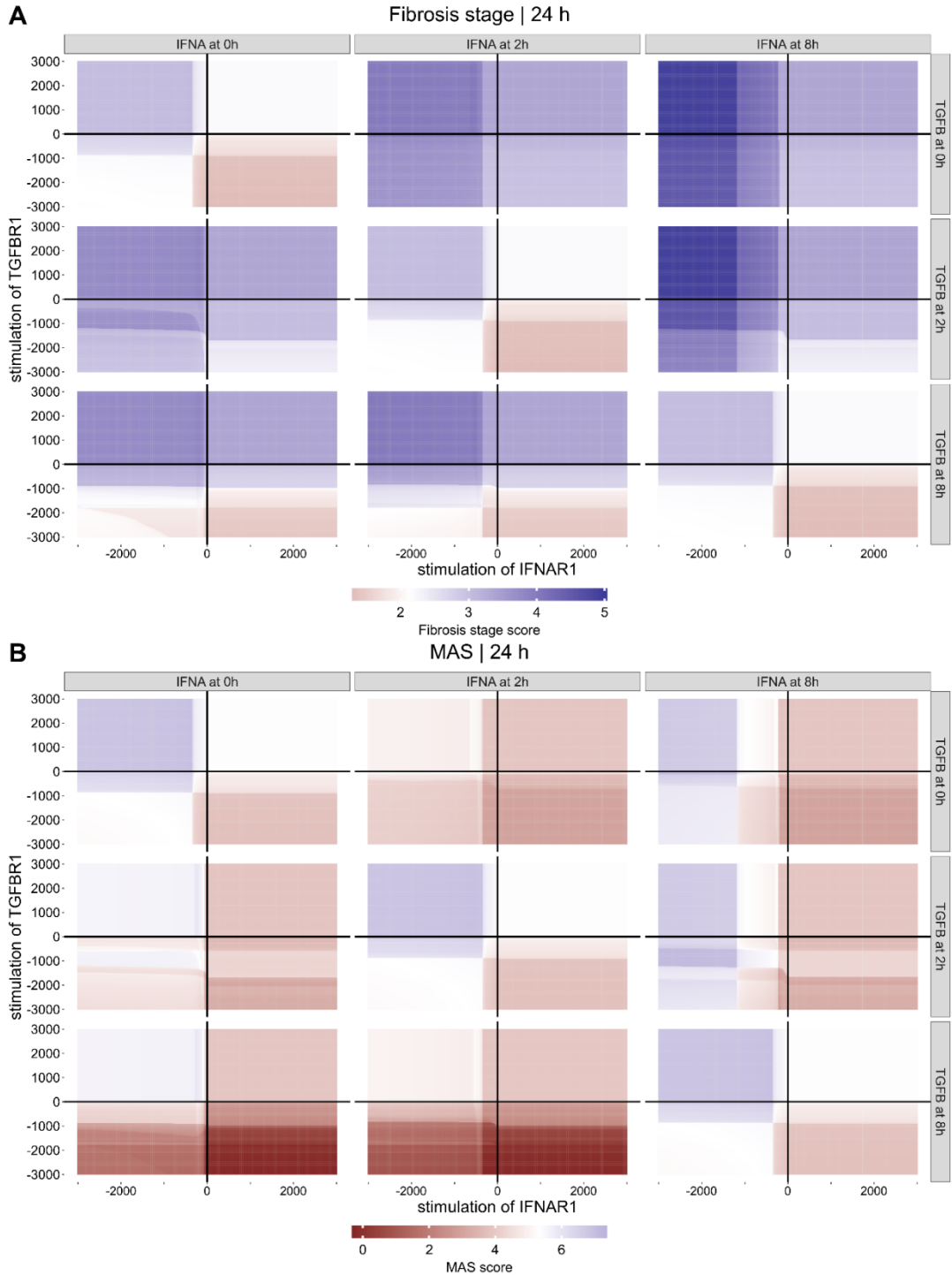

**Figure S10: Timing of TGFBR1 and IFNAR1 stimulation. A, B)** Ensemble-mean fibrosis stage (A) and MAS (B) scores measured at 24 hours. Rows and columns indicate TGFBR1 and IFNAR1 input onset at 0, 2, or 8 hours, respectively. Each heatmap spans receptor inputs from –3000 to +3000 arbitrary units. Color scales are centered on untreated scores.

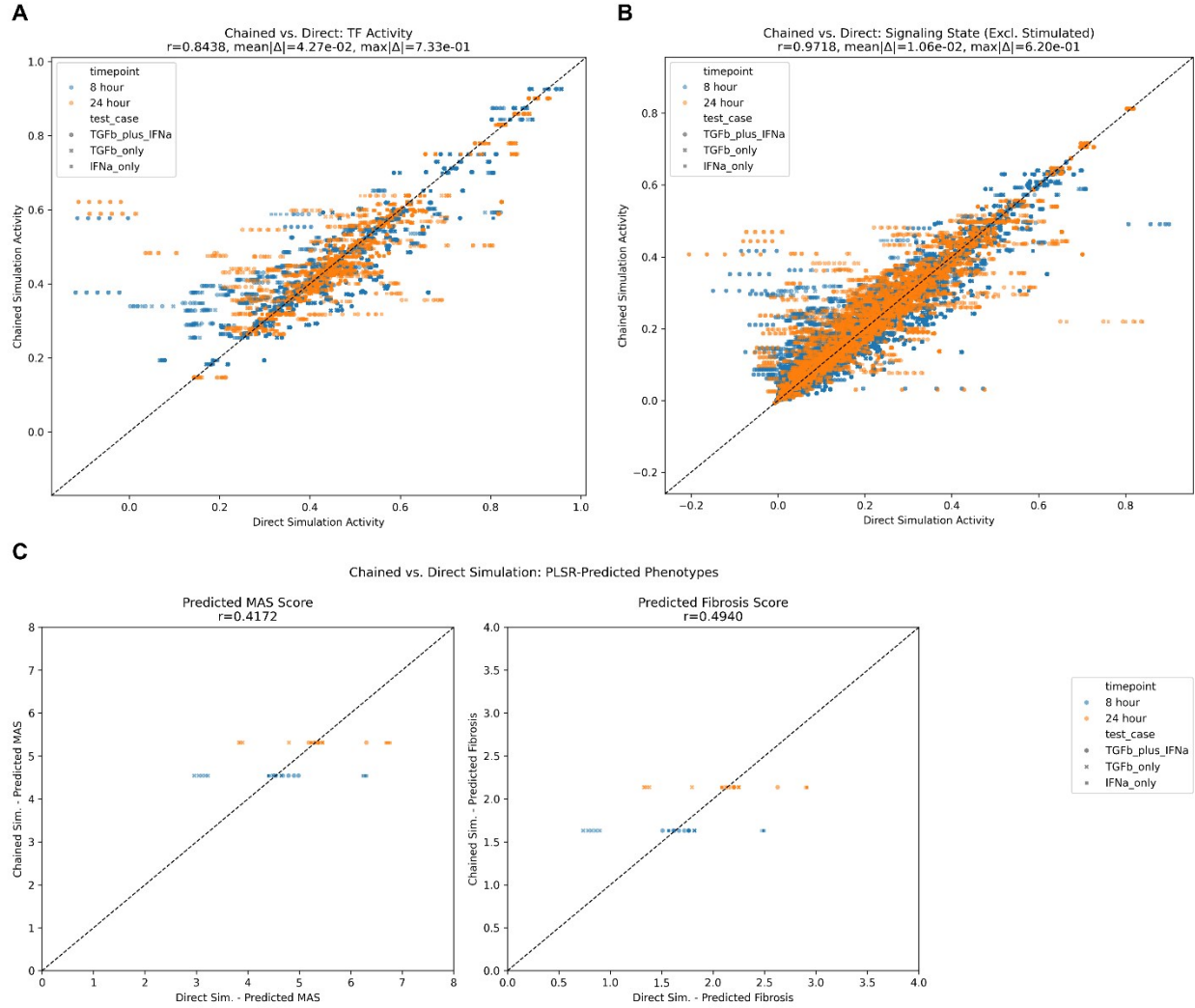

**Figure S11: Validation of sequential model-state carry-over. A–C)** Comparison of chained and direct simulations for TF activities (A), signaling node states excluding stimulated receptors (B), and PLSR-predicted MAS and fibrosis stage scores (C). Chained simulations transfer the ensemble-mean signaling state through the 2-, 8-, and 24-hour models. Colors indicate readout time; symbols indicate TGFBR1, IFNAR1, or combined stimulation. Dashed diagonals mark perfect agreement. $|\Delta|$ denotes absolute deviation between simulations.

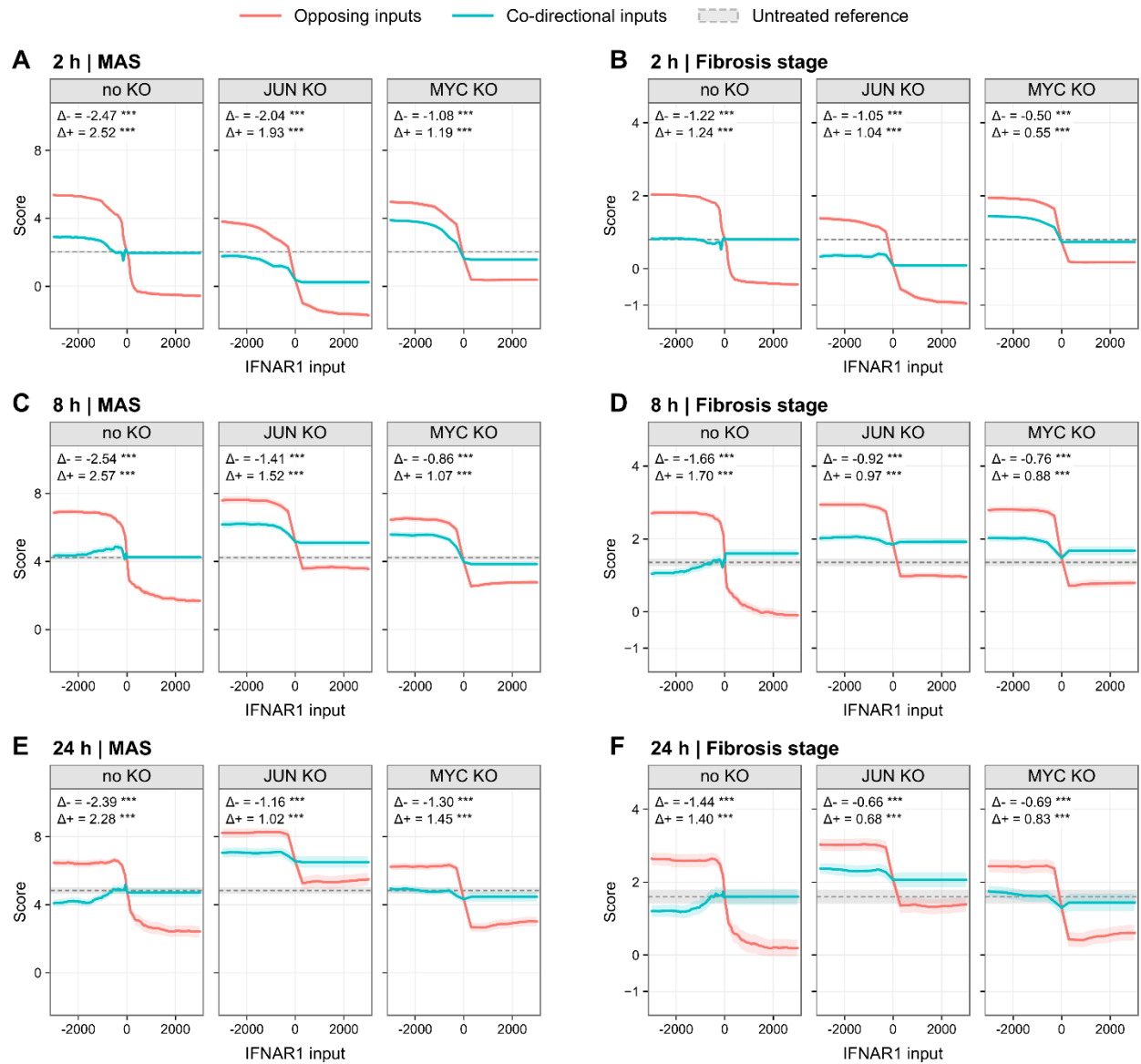

**Figure S12: JUN and MYC knockout during receptor co-stimulation. A–F)** Predicted MAS and fibrosis stage scores at 2 (A, B), 8 (C, D), and 24 (E, F) hours, without knockout or after JUN or MYC knockout. Coral and turquoise curves show opposing-sign and co-directional inputs, respectively. The horizontal axis denotes IFNAR1 input; TGFBR1 receives equal-magnitude input of opposite or matching sign. Lines show means across 50 models and colored ribbons $\pm 1$ SE. Grey shading shows untreated reference. $\Delta^-$ and $\Delta^+$ denote co-directional minus opposing-sign scores at inputs $-3000$ and $+3000$, respectively. Paired two-sided t-tests use Bonferroni adjustment within each phenotype and time point. \*\*\* p-value < 0.001.

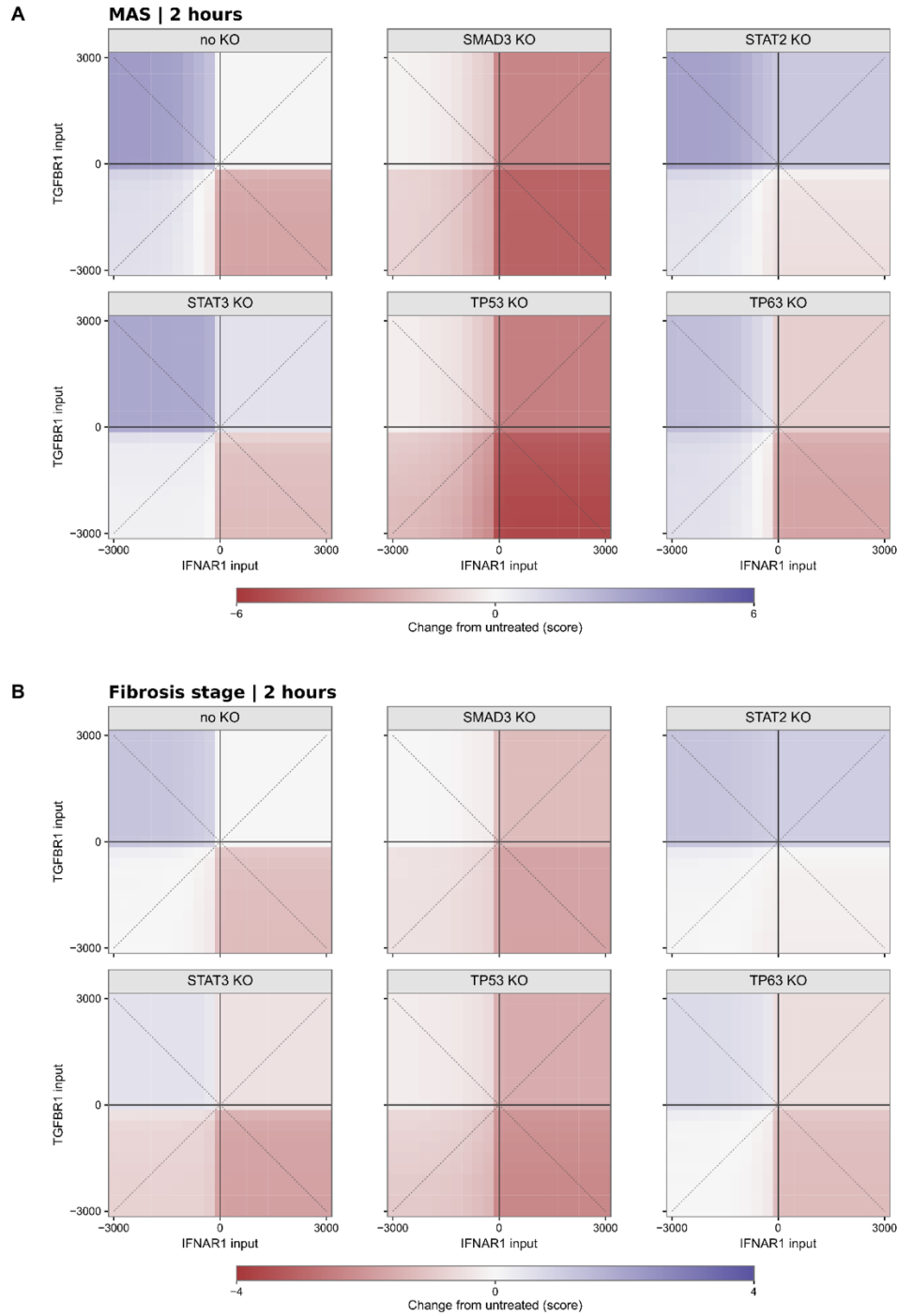

**Figure S13: Mediator knockouts across the receptor input grid at 2 hours. A, B)** Ensemble-mean changes in PLSR-predicted MAS (A) and fibrosis stage (B) scores without knockout or after SMAD3, STAT2, STAT3, TP53, or TP63 knockout, relative to the untreated no-knockout ensemble. Blue indicates increased scores and red indicates decreased scores. Black lines mark zero receptor input. Dashed diagonals mark opposing-sign and co-directional stimulation. Inputs are in arbitrary units; n = 50 models.

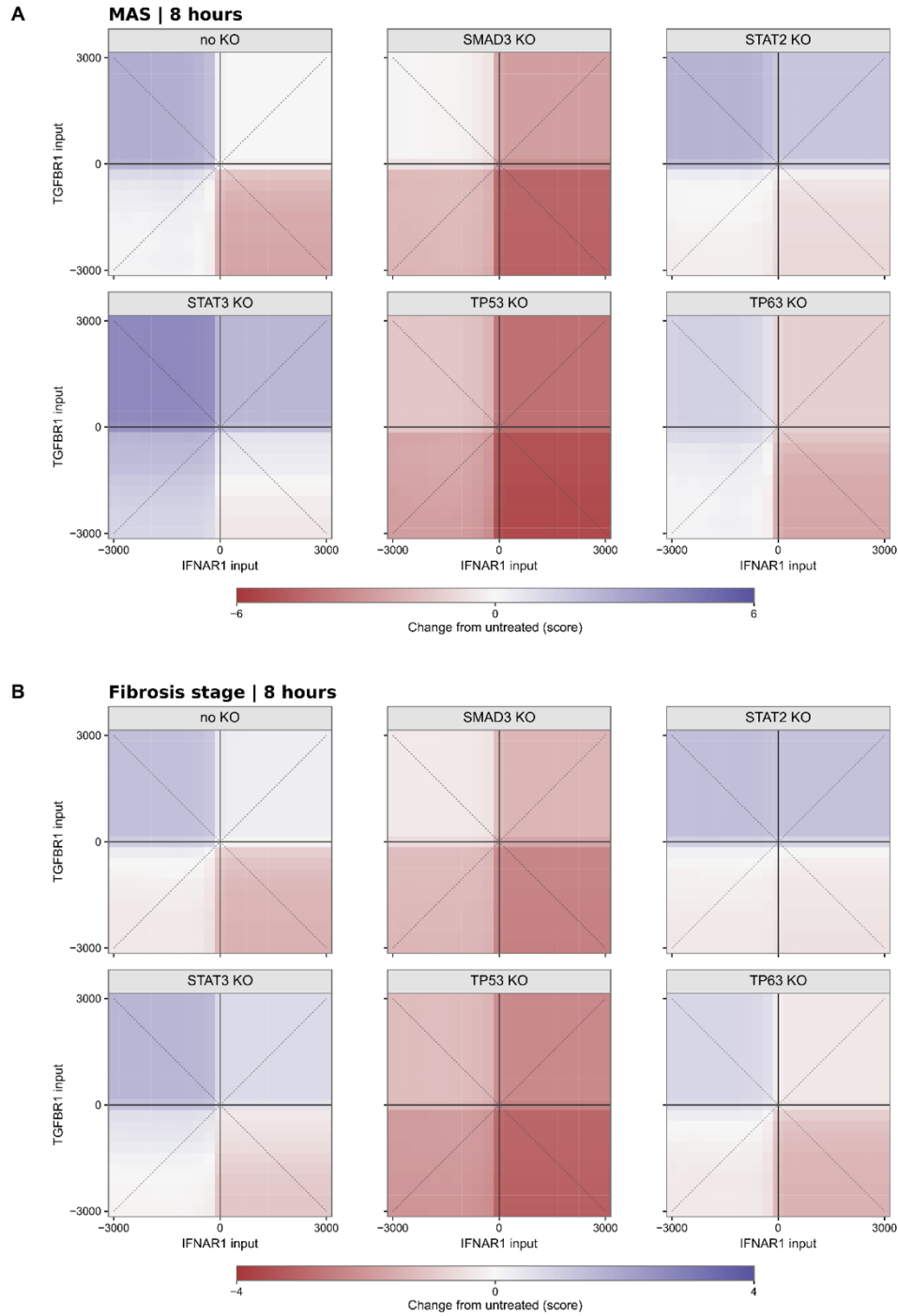

**Figure S14: Mediator knockouts across the receptor input grid at 8 hours. A, B)** Ensemble-mean changes in PLSR-predicted MAS (A) and fibrosis stage (B) scores without knockout or after SMAD3, STAT2, STAT3, TP53, or TP63 knockout, relative to the untreated no-knockout ensemble. Blue indicates increased scores and red indicates decreased scores. Black lines mark zero receptor input. Dashed diagonals mark opposing-sign and co-directional stimulation. Inputs are in arbitrary units; $n = 50$ models.

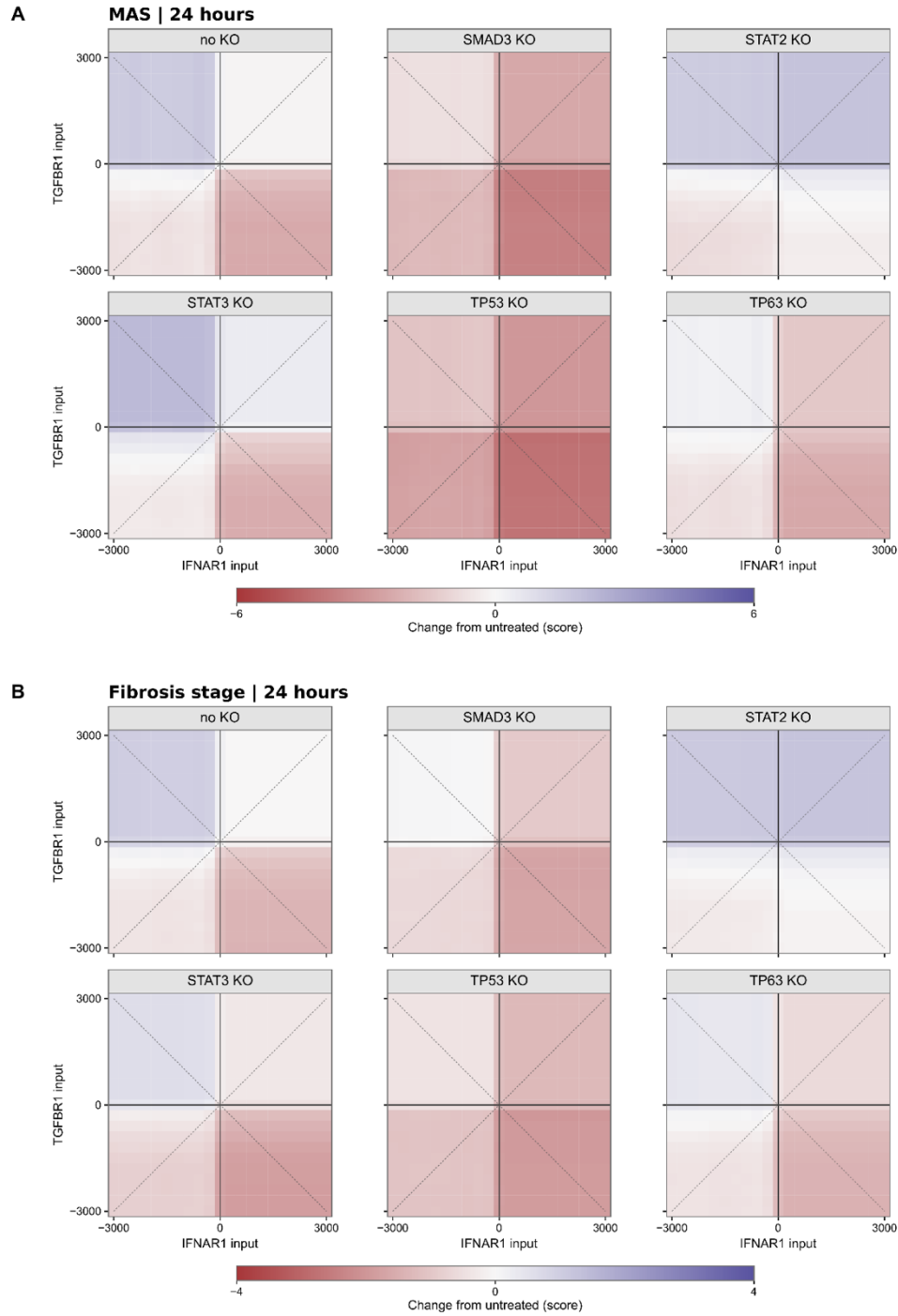

**Figure S15: Mediator knockouts across the receptor input grid at 24 hours. A, B)** Ensemble-mean changes in PLSR-predicted MAS (A) and fibrosis stage (B) scores without knockout or after SMAD3, STAT2, STAT3, TP53, or TP63 knockout, relative to the untreated no-knockout ensemble. Blue indicates increased scores and red indicates decreased scores. Black lines mark zero receptor input. Dashed diagonals mark opposing-sign and co-directional stimulation. Inputs are in arbitrary units; $n = 50$ models.

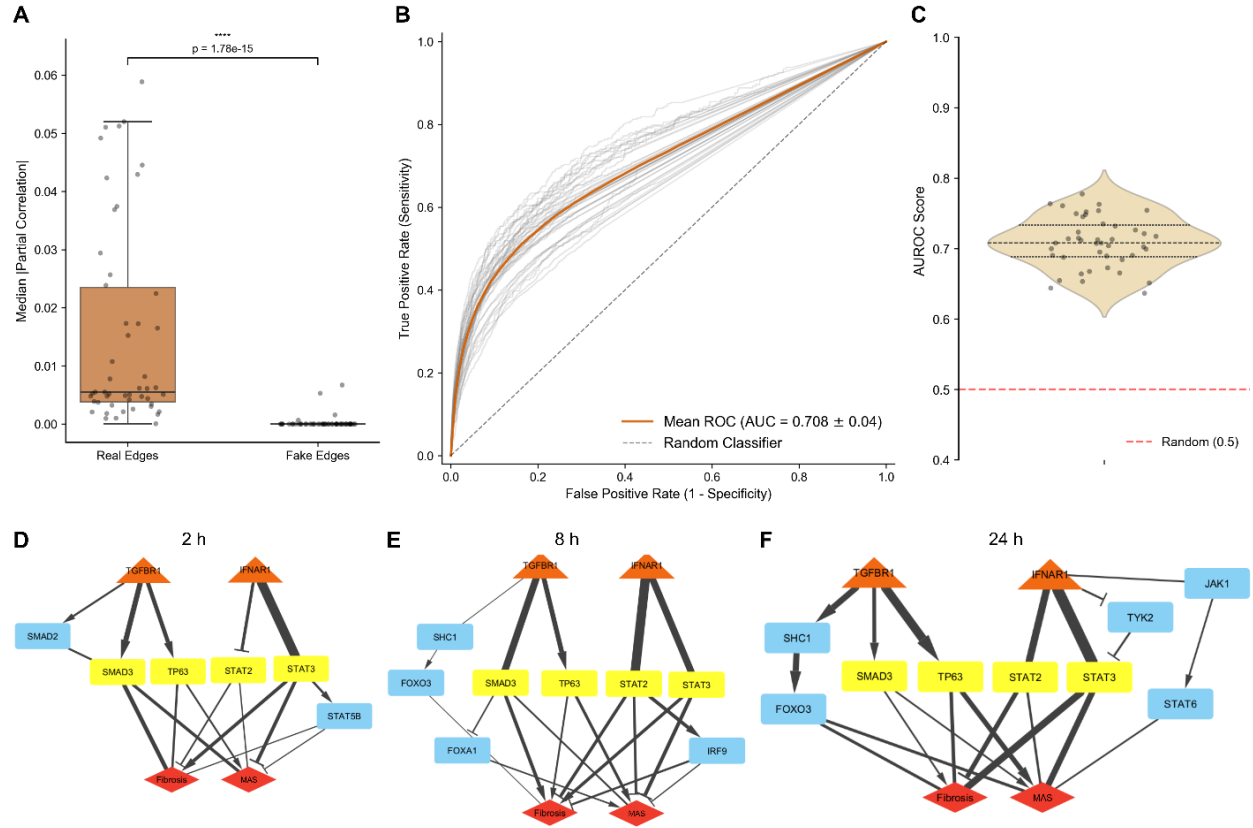

**Figure S16: Gaussian graphical model validation and PPS subnetworks.** **A)** Absolute partial correlation strengths for edges present (real) or absent (fake) in the signaling network, compared across 50 models using a paired Wilcoxon signed-rank test. **B, C)** ROC curves (**B**; grey, individual models; orange, mean) and model-level AUROC distribution (**C**). The diagonal in **B** and red line in **C** denote chance performance. **D–F)** Consensus networks from pair-path subscore (PPS) analysis at 2 (**D**), 8 (**E**), and 24 (**F**) hours. Yellow nodes highlight SMAD3, TP63, STAT2 and STAT3; line thickness reflects edge frequency among top paths. \*\*\*\*$p$-value $\leq 0.0001$.

### Supplementary Notes

#### **Supplementary Method 1: Sequential receptor stimulation and validation**

To investigate whether receptor stimulation order affects predicted phenotypes, we combined the ensembles of 50 DT-LEMBAS models trained separately at 2, 8, and 24 hours. TGFBR1 and IFNAR1 inputs were varied independently over 201 equally spaced values from –3000 to +3000 arbitrary units, yielding 40,401 combinations. Each receptor was stimulated at 0, 2, or 8 hours, giving nine protocols covering simultaneous stimulation and both sequential orders (Figure S10).

For sequential protocols, simulations started from a zero node/input vector. At each stage, the mean signaling node activities across the 50 models were passed as inputs to the next time point ensemble, together with any newly scheduled receptor stimulation. Receptor increments were added only at their scheduled times. For simultaneous protocols, the receptor/input vector was evaluated independently by each subsequent time point ensemble, without carrying forward the preceding output. Network propagation used up to 90 iterations, a leak parameter of 0.01, and a convergence tolerance of $10^{-5}$. Simulation timing was therefore represented through transitions between models trained at discrete time points.

Ensemble-mean transcription factor (TF) activities were clipped to [0, 1] and transformed using the stabilized logit, $\log[(\hat{x} + 10^{-6})/(1 - \hat{x} + 10^{-6})]$, where $\hat{x}$ denotes predicted TF activity. MAS and fibrosis stage scores were obtained using the same partial least-squares regression (PLSR) model described in the main Results and Methods of the study, fitted to clinical TF activities. Phenotype heatmaps show predictions at 24 hours, with color scales centered on the corresponding zero-stimulation reference.

Separately, we evaluated state transfer by comparing chained and direct simulations at 8 and 24 hours (Figure S11). TGFBR1 alone, IFNAR1 alone, or both receptors at equal input strengths were tested over 11 values spanning –3000 to +3000. In the chained validation procedure, the supplied vector initialized the recurrent node state, which was then updated using network weights and biases; ensemble-mean states were transferred through the 2-, 8-, and 24-hour models. Direct simulations applied the original receptor input vector to the corresponding time point ensemble. Agreement was quantified using Pearson correlation and mean and maximum absolute differences for TF activities and signaling node activities, excluding the two stimulated receptors from the latter comparison. Phenotype scores were compared using Pearson's correlation.

#### **Supplementary Method 2: Gaussian graphical modeling and pair-path subscore analysis**

To convert our signaling networks into a GGM, we utilized the Graphical Lasso (Glasso) algorithm implemented in Python (Schaipp *et al*, 2021). First, we utilized the same simulation grid that we utilized for the gradient-based decomposition method described in the main Results and Methods of the study. For each simulation, transcription factor activity scores were used to estimate a precision matrix. We then converted these into partial correlation matrices. The synthetic transcription factor data were first pre-processed to remove low-variance features (threshold = $10^{-3}$). Sample correlation matrices ($S$) were generated for each of the 50 ensemble simulations. Precision matrices ($\Theta$) were estimated using the gglasso library (Schaipp *et al*, 2021). Optimal sparsity parameters ($\lambda$) were selected using the extended Bayesian Information Criterion (eBIC) with $\gamma = 0.1$ (Chen & Chen, 2008). To mitigate shrinkage bias introduced by the $L_1$ penalty of Glasso, we refitted the estimated precision matrices using a Maximum Likelihood Estimation (MLE) constrained to the discovered sparsity pattern (Banerjee *et al*, 2008). The refitting was performed by minimizing the negative log-likelihood:

$$f(\theta) = -\log(\det(\theta)) + \text{tr}(S\theta),$$

using the L-BFGS-B optimization algorithm implemented in SciPy (Virtanen *et al*, 2020).

Refitted precision matrices were then converted into partial correlation matrices ($P$), where the partial correlation between nodes $i$ and $j$ is given by:

$$\rho_{ij} = -\frac{\theta_{ij}}{\sqrt{\theta_{ii}\theta_{jj}}}$$

To validate the performance of the model at differentiating between real edges and edges that did not exist, we calculated the Area Under the Curve (AUC) by comparing the inferred edge weights against a ground-truth network mask (Figure S16A-C). We also plotted the absolute value of the median partial correlation for both real edges and fake edges for each of the 50 ensemble models. Statistical significance was assessed using a pairwise Wilcoxon signed-rank test (Figure S16A).

We quantified the relative importance of specific signaling paths using the Pair-Path Subscore (PPS) algorithm (Gill *et al*, 2022). First, we added directionality to our GGM and removed noise by applying a mask containing the known edges and interactions in the DT-LEMBAS network to the partial correlation matrix. Then, for a specified source node $u$ (e.g., IFNAR or TGFBR) and target phenotype $v$ (e.g., fibrosis or NAS), we identified all simple paths $p$ up to a maximum length of $K = 5$ edges using a Depth-First Search (DFS) algorithm.

The contribution ($\eta_p$) of each path was calculated based on the path product ($\tau_p$) of partial correlations and the determinant of the submatrix $A$ (where $A_{ij} = -\rho_{ij}$ for $i \neq j$, and $A_{ii} = 1$) with the path's internal nodes removed:

$$\eta_p = \tau_p \cdot |A_{\setminus p}|$$

The final PPS ($s_p$) was defined as the absolute contribution of path $p$ normalized by the sum of absolute contributions of all paths between the pair:

$$s_p = \frac{|\eta_p|}{\sum_q |\eta_q|}$$

Top-ranking paths were aggregated across the 50 ensemble simulations to identify robust biological mechanisms (Figure S16D-F).
